# BRCA2-HSF2BP Oligomeric Ring Disassembly by BRME1 Promotes Homologous Recombination

**DOI:** 10.1101/2023.04.27.538421

**Authors:** Rania Ghouil, Simona Miron, Koichi Sato, Dejan Ristic, Sari E. van Rossum-Fikkert, Pierre Legrand, Malika Ouldali, Jean-Marie Winter, Virginie Ropars, Gabriel David, Ana-Andreea Arteni, Claire Wyman, Puck Knipscheer, Roland Kanaar, Alex N. Zelensky, Sophie Zinn-Justin

**Author notes:** These authors contributed equally (Rania Ghouil, Simona Miron).

## Abstract

In meiotic homologous recombination (HR), BRCA2 facilitates loading of the recombinases RAD51 and DMC1 at the sites of double-strand breaks. The HSF2BP-BRME1 complex interacts with BRCA2 to support its function in meiotic HR. In somatic cancer cells ectopically producing HSF2BP, DNA damage can trigger HSF2BP-dependent degradation of BRCA2, which prevents HR. Here we show that, upon binding to BRCA2, HSF2BP assembles into a large ring-shaped 24-mer consisting of three interlocked octameric rings. Addition of BRME1 leads to dissociation of this ring structure, and cancels the disruptive effect of HSF2BP on cancer cell resistance to DNA damage. It also prevents BRCA2 degradation during inter-strand DNA crosslink repair in *Xenopus* egg extracts. We propose that the control of HSF2BP-BRCA2 oligomerization by BRME1 ensures timely assembly of the ring complex that concentrates BRCA2 and controls its turnover, thus promoting meiotic HR.

## INTRODUCTION

In vertebrates, both somatic and meiotic homologous recombination (HR) require the BRCA2 protein^1, 2^. Its orthologues in fungi^3^, plants^4^ and invertebrates^5–8^ are also essential for meiotic HR, so this role is likely ancestral. Most mechanistic data on BRCA2, however, comes from studies in somatic cells, due to the strong association of BRCA2 with breast, ovarian, pancreatic and some other types of cancer^9^. These studies showed that BRCA2 interacts with and controls RAD51, a recombinase that performs the central HR reactions: homology recognition and strand exchange. Meiotic studies are hindered by the embryonic lethality of the *Brca2* knock-out in mice. A hypomorphic rescue transgene in a *Brca2* knock-out mouse strain confirmed the critical role of BRCA2 in mouse meiotic HR^10^. It was also demonstrated that BRCA2 interacts with the meiotic recombinase DMC1^11–14^, a paralogue of RAD51. This suggested that BRCA2 may contribute to the correct balance in DMC1 and RAD51 loading that is essential in meiotic HR.

New tools to study the role of BRCA2 in meiosis were recently provided by the discovery that in mouse meiocytes and embryonic stem cells BRCA2 functions in complex with two previously uncharacterized germline proteins, HSF2BP (also called MEILB2) and BRME1 ^15–18^. The reported phenotypes of three independent *Hsf2bp*^17–19^ and five independent *Brme1* knock-out mouse models^16–22^ are nearly identical. Loss of these proteins does not lead to embryonic lethality, but causes complete spermatogenesis failure due to severe reduction in RAD51 and DMC1 accumulation at meiotic DNA double strand breaks (DSBs), which prevents crossover formation^17, 18^. Loss of BRME1 decreases the number of HSF2BP foci. As HSF2BP directly binds to both BRCA2 and BRME1 (Fig. 1a), it has been proposed that loss of HSF2BP or BRME1 compromises BRCA2 meiotic function, although the mechanism may be different ^23^ from the suggested “meiotic localizer of BRCA2” model ^18^.

**Fig. 1.**
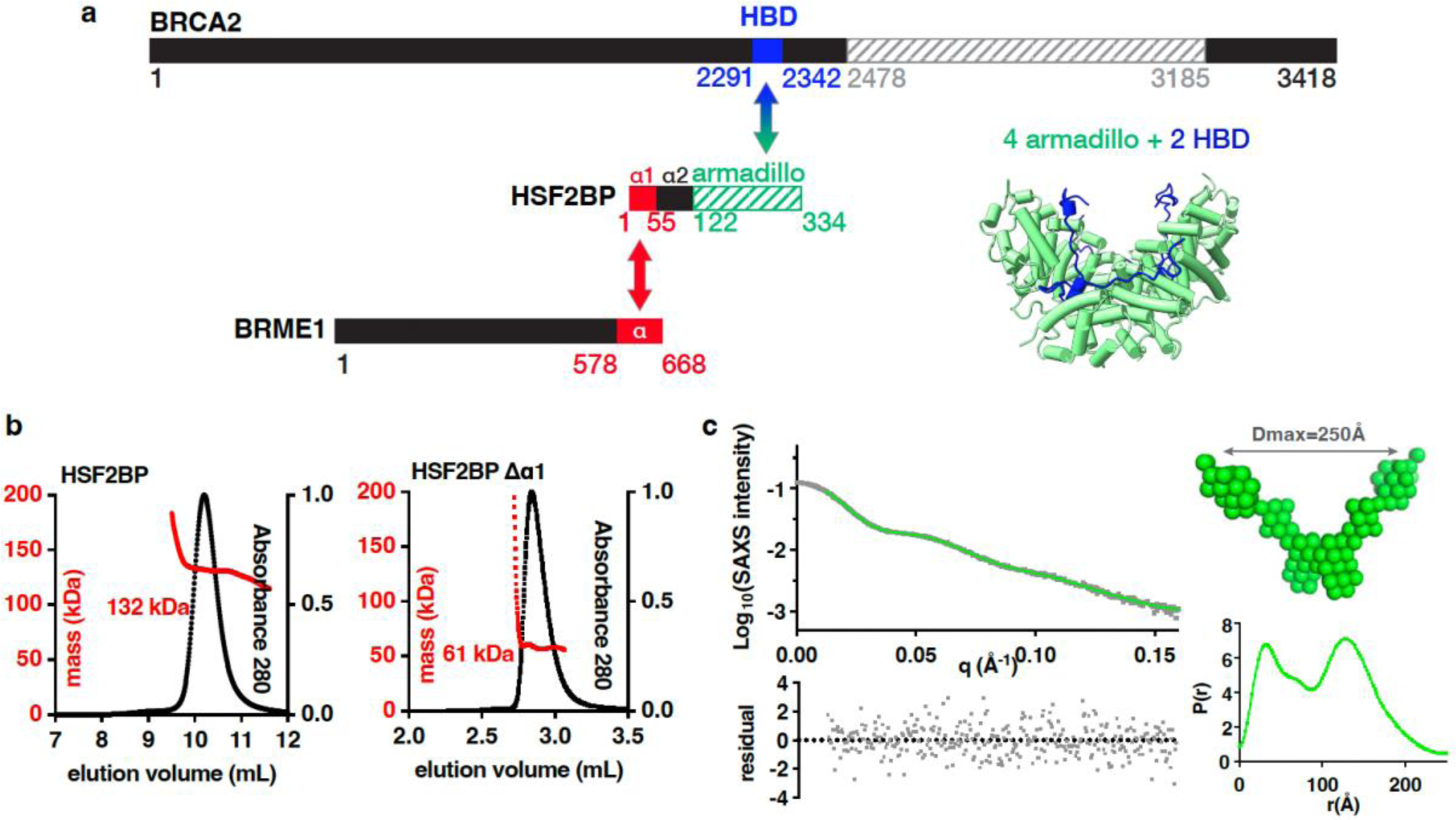
HSF2BP is a tetramer formed by two dimers interacting through the N-terminal helix α1. **a**, BRCA2, HSF2BP and BRME1 domains and interactions. Folded domains of known 3D structure are marked with stripes. Co-crystallized regions of BRCA2 (HBD) and HSF2BP (armadillo) are colored in blue and green, respectively; the 3D structure cartoon is shown (PDB code: 7BDX). Regions of HSF2BP and BRME1 that interact are colored in red ^16^. **b**, SEC-MALS analysis of full-length HSF2BP (monomer Mw 37.6 kDa) and HSF2BP deleted from helix α1 (monomer Mw 32.2 kDa). See Extended Data Fig. 1a for replicates. **c**, SEC-SAXS curve and resulting distance distribution, obtained on full-length HSF2BP. The SAXS curve is plotted as a function of the scattering angle (experimental curve in black dots; P(r) Fourier Transform in green). The distance distribution P(r) is plotted as a function of the distance. The deduced HSF2BP mass is 156 ± 21 kDa. Calculation of an *ab initio* model (average model in green spheres; more models in Extended Data Fig. 1b,c) from the SAXS data suggests that HSF2BP has a V shape.

Paradoxically, when HSF2BP is produced ectopically in somatic cancer cells, it suppresses HR instead of supporting it as it does in meiocytes^15, 24^. We demonstrated genetically and biochemically that this is due to HSF2BP interaction with BRCA2: in these cells, formation of a complex between HSF2BP and BRCA2 impedes BRCA2 function during the repair of lesions induced by DNA interstrand crosslinking agents and PARP inhibitors (but not by ionizing radiation or the I-SceI nuclease) ^15, 24^. Evolutionary conservation of the interaction between BRCA2 and HSF2BP allowed us to establish, using *Xenopus* egg extract interstrand crosslink repair assays, that the presence of HSF2BP leads to DNA crosslink-dependent proteasomal degradation of BRCA2 mediated by the p97 segregase ^24^, which disassembles ubiquitinated protein aggregates ^25, 26^.

To study the molecular events triggered by the interaction between BRCA2 and HSF2BP, we previously solved the crystal structure of a 51-aa BRCA2 peptide (BRCA2-HBD) in complex with the C-terminal armadillo domain of HSF2BP (Fig. 1a). This revealed a cryptic motif repeated twice in BRCA2-HBD and encoded by exons 12 and 13, respectively. Each motif binds to one armadillo domain ^23, 27^. In the complex, two BRCA2-HBD peptides “staple together” two armadillo dimers, resulting in a high affinity interaction (Kd ∼1 nM). In addition to the crystallized armadillo domain, HSF2BP contains N-terminal helices α1 and α2 (Fig. 1a) that assemble into oligomers in vitro. Helix α1 homotetramerizes when free in solution, and heterotetramerizes when bound to the C-terminal α-helical region of BRME1 ^18, 19, 21^. Presence of these three HSF2BP oligomerization mechanisms — dimerization via armadillo and α2, tetramerization via α1 and hetero-oligomerization via BRCA2-HBD binding — led us to propose that HSF2BP is a polymerization agent for BRCA2, which could be behind its physiological function in meiotic HR and mediate its pathological effect in somatic HR ^23^. However, since the N-terminal helices of HSF2BP were missing from previous structural analyses, their effect on the HSF2BP-BRCA2 structure and function was uncharacterized ^23^. It was also not known how the interaction between HSF2BP helix α1 and BRME1 affects the oligomeric state of the BRCA2-HSF2BP complex and BRCA2 function in cells.

In this work, we obtained a cryo-EM derived model of the complex between full-length HSF2BP and BRCA2-HBD, determined how BRME1 affects the structure of this complex, and established a model explaining the opposite effects of HSF2BP on somatic and meiotic HR. Our work revealed that, upon binding to BRCA2, HSF2BP forms a large (⌀∼200Å) ring-shaped 24-mer consisting of three interlocked octameric rings, with BRCA2 displayed on the outer surface of the ring. We also demonstrated that BRME1 disrupts this ring structure and acts as a protective disaggregation agent in specific cellular conditions.

## RESULTS

### HSF2BP is mainly a V-shaped tetramer in solution

To understand how the N-terminal α-helical region of HSF2BP contributes to the assembly of the HSF2BP-BRCA2 complex, we first characterized full-length HSF2BP (37.6 kDa) and HSF2BP lacking helix α1 (G48-V334; 32.2 kDa) by size-exclusion chromatography coupled to multi-angle light scattering (SEC-MALS). At the concentration of the experiment (∼10-30 μM), the apparent molar masses of the two samples are 136 ± 7 kDa (n=2) and 60 ± 2 kDa (n=2), respectively (Fig. 1b; Extended Data Fig. 1a). This fits with tetrameric and dimeric states for HSF2BP with and without helix α1, respectively. Further characterization of HSF2BP by size-exclusion chromatography coupled to small-angle X-ray scattering (SEC-SAXS) showed that, in the conditions of this experiment (protein concentration ∼50-60 μM), the protein has a molar mass of 156 ± 21 kDa (Fig. 1c). Thus, we confirmed that full-length HSF2BP is mainly tetrameric at concentrations above 10 μM. The SAXS-derived atomic distance distribution curve of HSF2BP is bimodal, with a maximal distance at 250 Å (Fig. 1c). Consistently, *ab initio* molecular envelopes calculated from this curve using a 2-fold symmetry hypothesis have a V shape (Fig. 1c; Extended Data Fig. 1b,c) ^21^. Altogether, our data support a full-length HSF2BP model in which two dimeric fragments containing helices α2 and armadillo domains are connected through the tetrameric N-terminal helix α1.

### HSF2BP and BRCA2 form a ring-shaped complex with BRCA2 on its outer surface

Having characterized the solution structure of free full-length HSF2BP, we measured the change in its oligomeric state induced by BRCA2. Tetrameric HSF2BP forms an 880 ± 30 kDa (n=2) complex when bound to BRCA2-HBD (N2291-Q2342), as measured by SEC-MALS (Fig. 2a; Extended Data Fig. 1a). Its thermal stability also increases substantially, shifting from 45.9-46.7°C to 59-60.7°C (Fig. 2b). Negative-staining electron microscopy (EM) revealed a ring-shaped complex with an outer diameter of about 200 Å (Fig. 2c) and a 3-fold symmetry (Extended Data Fig. 2a). Further characterization by cryo-EM (Extended Data Fig. 2b,c) revealed a *D3* symmetry, defined as a 3-fold symmetry around the axis going through the center of the ring, as well as three 2-fold symmetries around perpendicular axes going through the center of the globular subvolumes (Fig. 2d). The most resolved cryo-EM map was calculated from 398,000 particles and refined using this symmetry. Resulting local resolutions yield 3-4.5 Å and 3.5-5 Å in the three inner and outer globular subvolumes, respectively. The tubular-shaped volumes extending out of the globular subvolumes (two tubes on both sides of each subvolume) are less resolved (4-8 Å). Further 3D classification of the cryo-EM particles resulted in the calculation of two cryo-EM maps. Comparison of these maps revealed that the diameter of the ring is slightly variable, with the larger ring being generally better resolved than the tighter ring, except for the inner globular subvolumes that are better resolved in the tighter ring (Extended Data Fig. 2d). The heterogeneous resolution of the calculated cryo-EM maps might therefore result from the ring flexibility. We fitted the published crystal structure of the C-terminal armadillo domain of HSF2BP in complex with BRCA2-HBD^23, 27^ into each of the six globular subvolumes of the ring (Fig. 2e). Remarkably, not only the α-helices of the armadillo domains, but also the BRCA2-HBD peptides can be traced through the map (Fig. 2f; Extended Data Fig. 2c); several side chains can even be identified. The BRCA2 peptides are located on the outer side of the map, their N- and C-termini being positioned around the ring complex (Fig. 2g).

**Fig 2.**
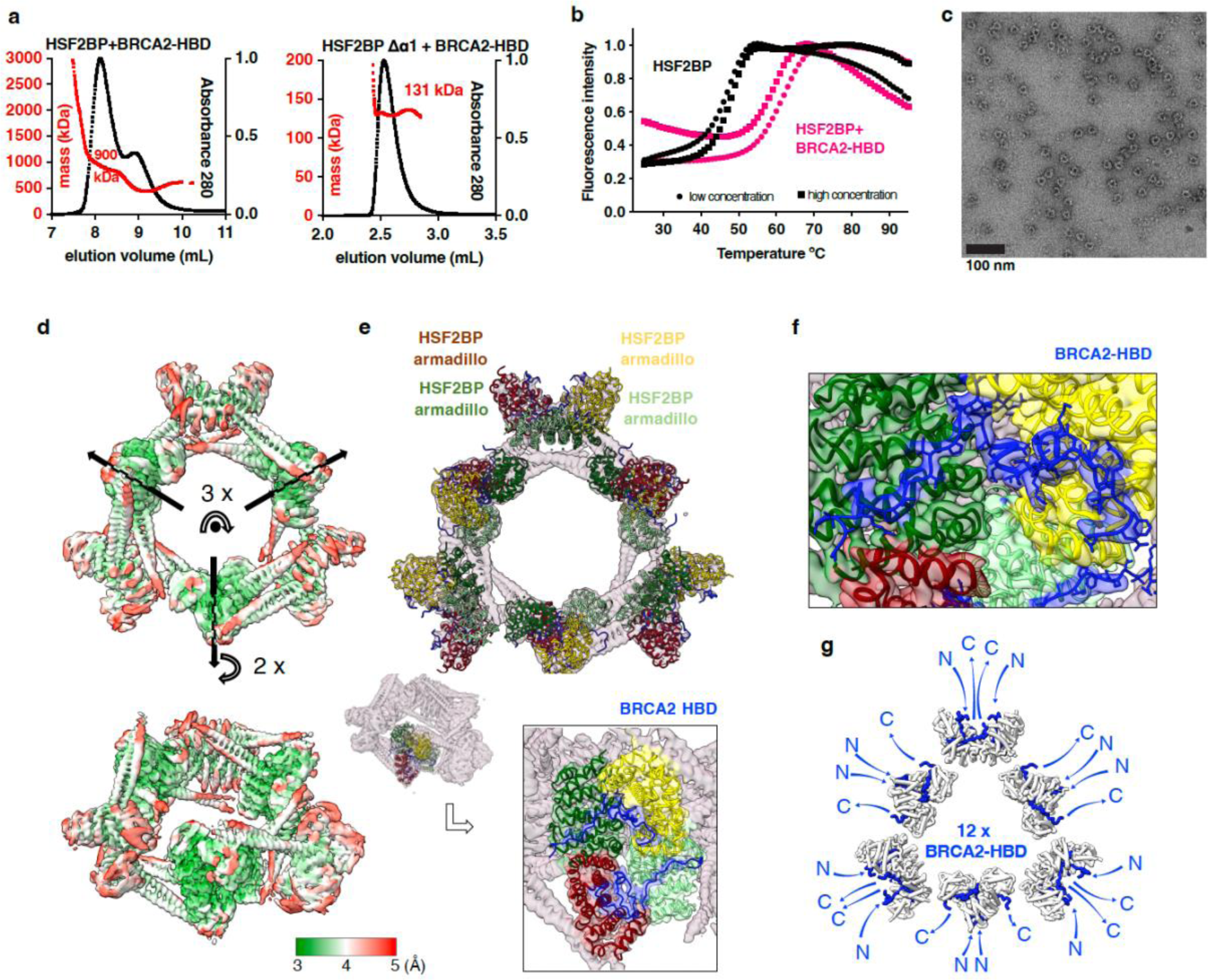
HSF2BP oligomerizes into a ring-shaped complex upon binding to BRCA2-HBD. **a**, SEC-MALS analysis of HSF2BP (either full length or deleted from helix α1) bound to BRCA2-HBD. See Extended Data Fig. 1a for replicates. **b**, Thermal stability of HSF2BP, either free or bound to BRCA2-HBD. The denaturation temperature of HSF2BP shifted from 45.9-46.7°C to 59-60.7°C upon binding to BRCA2-HBD. **c**, Negative-staining EM image obtained on a sample of HSF2BP bound to BRCA2-HBD. **d**, Cryo-EM map of HSF2BP bound to BRCA2-HBD. Top and side views are displayed with an electron density threshold of 0.05. The map shows a *D3* symmetry. It is colored as a function of the local resolution: from green (3 Å) to red (5 Å). **e**, Docking of the crystal structure of the complex between HSF2BP armadillo domain and BRCA2-HBD (PDB: 7BDX) into the cryo-EM map. In the top view, a crystal structure is positioned in each of the six globular subvolumes of the map, whereas in the side view, only one crystal structure is displayed. Each crystal structure contains four armadillo domains (in light green, yellow, green and maroon) bound to two BRCA2-HBD peptides (in blue). In the boxed side view, the map is colored as the docked chains of the crystal structure. **f**, Zoom view on one of the BRCA2 peptides docked into the cryo-EM map. The position of this peptide, represented in blue sticks, readily fits into the cryo-EM density map of the complex between HSF2BP and BRCA2-HBD. The map is colored as the docked chains of the crystal structure. **g,** Orientations of the N- and C-termini of the 12 BRCA2 peptides in the complex. All the BRCA2 extremities are located on the outer surface of the ring shape.

### HSF2BP helices α1 and α2 connect BRCA2-bound HSF2BP armadillo tetramers in the HSF2BP-BRCA2 complex

We further analyzed our cryo-EM map of the complex between HSF2BP and BRCA2-HBD, in order to describe the 3D structure of HSF2BP N-terminal α-helical region in this complex. First, we modelled a dimer of HSF2BP using AlphaFold ^28^. This prediction algorithm proposed five similar models for dimeric HSF2BP with high confidence (Extended Data Fig. 3a). In these models, each HSF2BP monomer is composed of strand β1 (F20-R24), helix α1 (K25-I45) and helix α2 (G48-S136) that overlaps with the armadillo domain (E122-V334). As a validation, we checked that the 3D model of the dimeric armadillo domain is superimposable with the 3D structure observed in the crystal (PDB code: 7BDX ^23^, Extended Data Fig. 3b). We then docked two dimeric HSF2BP models into the cryo-EM map of the complex between full-length HSF2BP and BRCA2-HBD (Fig. 3a). Helices α2 readily fitted into the electron density tubes extending out of the globular subvolumes corresponding to the armadillo domains bound to BRCA2-HBD. Helices α1, on the other hand, fell into a poorly resolved zone connecting electron densities of two HSF2BP dimers (Fig. 3a,b). In the resulting model of the complex, two HSF2BP dimers interact through their helices α1 to form a V-shape tetramer. These tetramers interact two by two through the BRCA2-HBD peptides to form octamers. Three interlocked octamers (displayed in red, blue and green in Fig. 3a, inset) form the final ring-shaped 24-mer protein assembly. The N-terminal region of HSF2BP contributes to the assembly of this large complex through interactions between helices α1 within each tetramer and contacts between helices α2 and armadillo domains from different octamers (Fig. 3b).

**Fig. 3.**
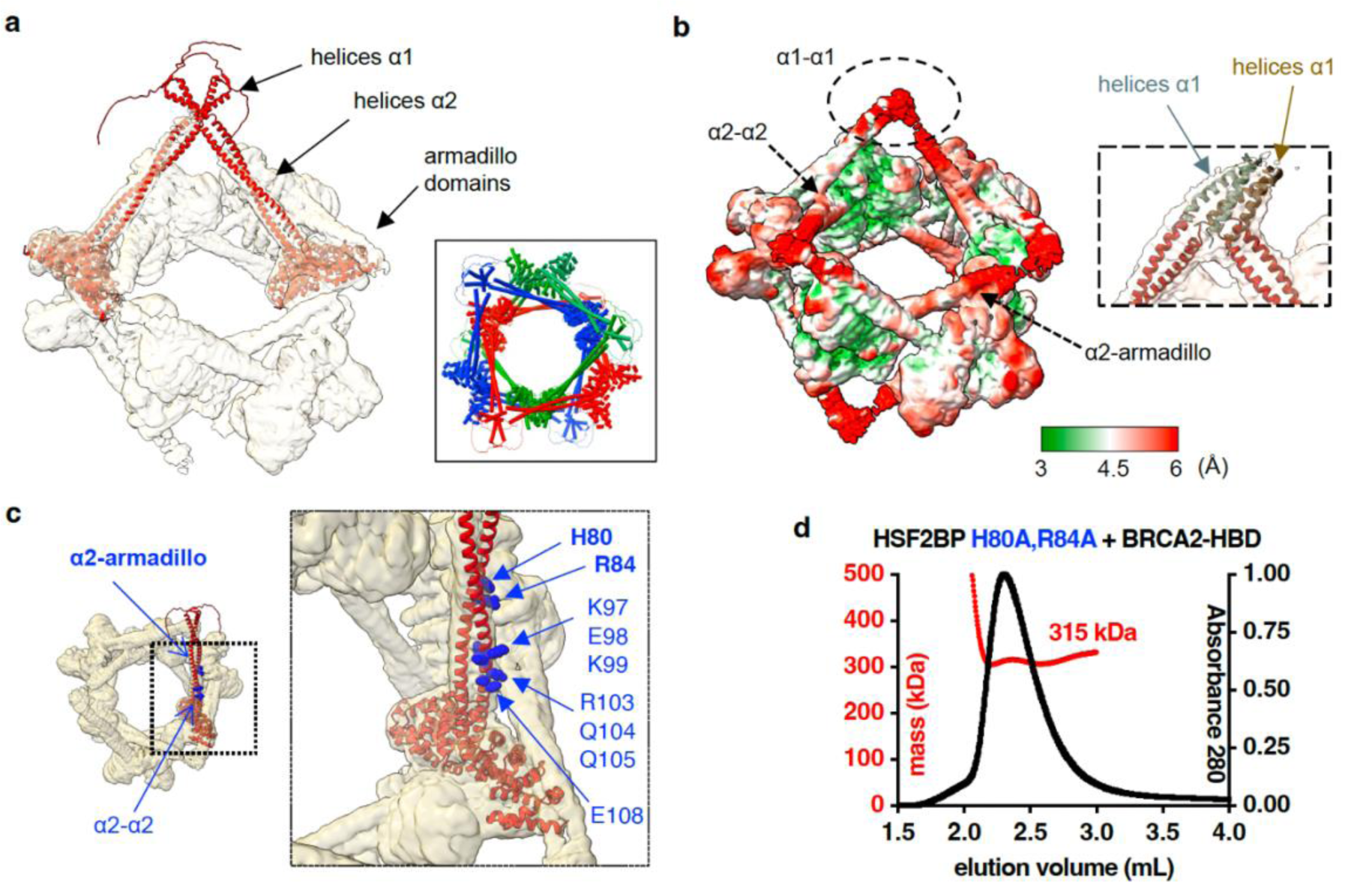
The ring-shaped complex is assembled through a large set of interfaces, involving not only armadillo-BRCA2 but also α1-α1, α2-α2 and α2-armadillo contacts. **a**, Docking of two (similar) AlphaFold models of the HSF2BP dimer into the cryo-EM map. The map is displayed with an electron density threshold of 0.035. The cartoon views of the two AlphaFold models are colored in red. Each model consists of a disordered region, a short helix α1, a large helix α2 and an armadillo domain. In the boxed panel, 3 pairs of HSF2BP tetramers docked into the cryo-EM map are displayed in 3 different colors**. b**, Cryo-EM map displayed with a lower electron density threshold (0.025) and colored as a function of the local resolution. The colors are from green (3 Å) to red (6 Å). A dashed oval identifies a map region corresponding to the 4 helices α1, whereas arrows indicate regions of α2-α2 and α2-armadillo contacts. In the inset, 4 helices α2 are docked as in a) (in red), whereas 4 parallel helices α1 are manually positioned into the cryo-EM density (in grey and brown). **c**, Position of the α2 residues mutated to test their role in the assembly and/or function of the ring-shaped complex. An AlphaFold model of HSF2BP dimer (in red) is docked into the cryo-EM map. Residues at the α2-α2 and α2-armadillo interfaces are marked with standard and bold labels, respectively. **d**, SEC-MALS analysis of the mutant HSF2BP H80A-R84A bound to BRCA2-HBD (see also Extended Data Fig. 4).

To further understand how the N-terminal region of HSF2BP contributes to the assembly of this large complex, we modelled the α1 tetramer using AlphaFold (Extended Data Fig. 3c). The best scored model is composed of two anti-parallel α1 dimers that cannot be fitted in the density map. However, several parallel tetramers were also proposed by AlphaFold (models 3-5). They are similar to each other and closer to what can be obtained by fitting four α1-helices into the cryo-EM density map (Fig. 3b, inset). The low resolution of this part of the map is probably due to the variable position of the tetramer relatively to the ring; it prevents a more detailed description of the 3D structure of this tetramer.

We also analyzed how the octamers interact to form the large ring-shaped complex. The three diamond-shaped octameric rings (Fig. 3a, inset) could exist as separate complexes, yet they interlock to form the bigger 24-mer complex. This suggests that a network of contacts between octamers favors the formation and high thermal stability of the 24-mer complex and that a dynamic mechanism exists for octamer opening to allow interlocking. Two types of contacts are observed between octamers, involving either coiled coils of helices α2 or one of these coiled coils and an armadillo domain (indicated by arrows in Fig. 3b,c). In each octamer, two α2 coiled coils interact with a second octamer (specifically, with its helices α2 and armadillo domains), whereas the two other α2 coiled coils interact with the third octamer (also with its helices α2 and armadillo domains). We engineered mutations in HSF2BP helix α2, in order to specifically disrupt these contacts. We mutated a set of selected residues into alanine (Fig. 3c), and measured the oligomeric state of the resulting free and BRCA2-bound HSF2BP variants by SEC-MALS (Fig. 3d; Extended Data Fig. 4). Remarkably, the H80A-R84A variant only assembled into octamers upon binding to BRCA2-HBD; it formed no 24-mers (Fig. 3d). Thus, not only the α1-α1 interaction but also the α2-armadillo interaction contribute to the formation of the large BRCA2-bound HSF2BP oligomer.

### HSF2BP helix α1 forms a heterotetramer with an α-helical peptide from the C-terminal region of BRME1

Mouse HSF2BP binds directly to the meiotic protein BRME1 ^16^. The N-terminal helix α1 of HSF2BP interacts with the C-terminal region of BRME1 ^16, 20^. To further delineate the binding fragment in human BRME1, we divided its C-terminal region (D578-L668) into three peptides: BRME1-M, corresponding to the central and well-conserved fragment, and BRME1-N and BRME1-C corresponding to the less conserved flanking regions (Fig. 4a). Isothermal titration calorimetry (ITC) experiments revealed that only BRME1-M significantly bound to HSF2BP, and that its affinity was similar when measured against full-length HSF2BP (18 ± 3 nM) or HSF2BP helix α1 (25 ± 5 nM) (Fig. 4b, Extended Data Fig. 5a, Table 1). Thus, the C-terminal region BRME1-M (E602-K641) directly interacts with the HSF2BP N-terminal helix α1. This interaction is not expected to affect binding of BRCA2 to the HSF2BP C-terminal armadillo domain^23^. Consistently, we found by ITC that BRME1-M is able to interact with HSF2BP bound to BRCA2–HBD (affinity: 34 ± 8 nM), and that BRCA2-HBD is able to interact with HSF2BP bound to BRME1-M (affinity: 13 ± 2 nM) (Fig. 4c). We concluded that HSF2BP simultaneously interacts with BRCA2-HBD and BRME1-M.

**Fig. 4.**
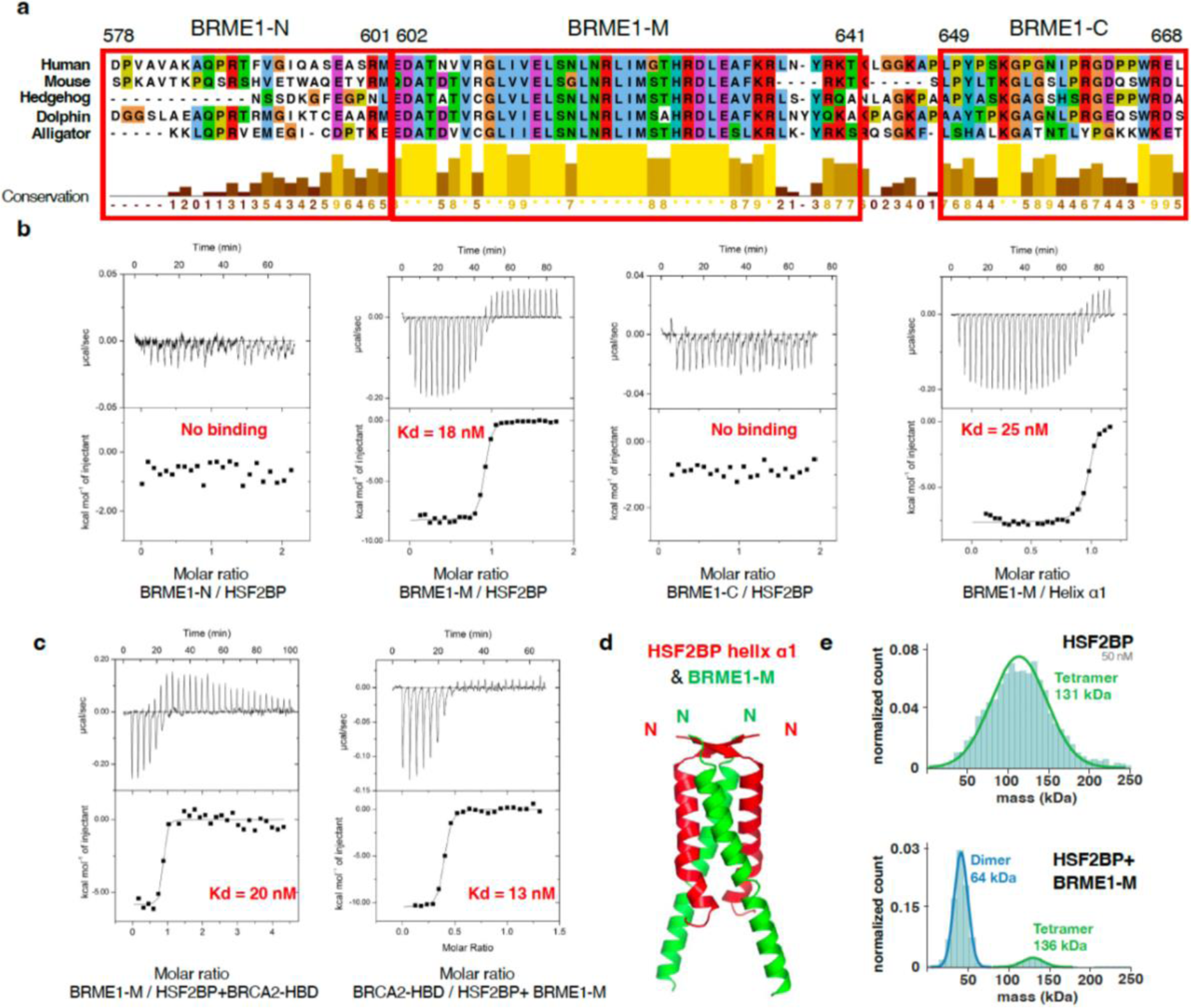
A BRME1 peptide binds to helix α1 and disrupts the HSF2BP tetramer. **a**, Sequence alignment of the C-terminal region of a set of five BRME1 proteins, showing a representative sequence diversity. The sequences corresponding to the three human peptides BRME1-N, BRME1-M and BRME1-C are boxed. **b**, ITC curves identifying the BRME1 sequence binding to HSF2BP. These experiments were all performed at 30°C. Additional experiments performed at 20°C are detailed in Extended Data Fig. 5a and Table 1. **c**, ITC curve showing that BRME1-M and BRCA2-HBD do not compete for binding to HSF2BP. These experiments were performed at 20°C. **d**, Crystal structure of the HSF2BP peptide E19-V50 (helix α1, in red) bound to the BRME1 peptide E602-K641 (BRME1-M, in green). Each asymmetric unit contained a HSF2BP-BRME1 dimer. The heterotetramer was calculated by application of a 2-fold crystallography symmetry (Supplementary Table 1). **e**, Mass photometry experiment performed on either free HSF2BP (upper panel) or HSF2BP bound to BRME1-M (lower panel). The distribution of masses is displayed for samples at a concentration of 50 nM for HSF2BP and 100 nM for BRME1-M. HSF2BP analyzed at 25 nM is shown in Extended Data Fig. 6c.

**Table 1.** Binding parameters deduced from the ITC analyses.

| | Kd (M) (± error) | n | $\Delta H$ (kcal/mol)<br>(± error) | $\Delta G$<br>(kcal/mol) | $-T\Delta S$<br>(kcal/mol) | T (K) |
| --- | --- | --- | --- | --- | --- | --- |
| HSF2BP vs BRME1-M | 1.1E-08 (3.5) | 0.77 | -5.7 (0.07) | -10.6 | -4.90 | 293 |
| HSF2BP vs BRME1-M | 1.8E-08 (2.7) | 0.88 | -8.2 (0.06) | -10.7 | -2.50 | 303 |
| HSF2BP vs BRME1-N | undetectable |  |  |  |  | 293 |
| HSF2BP vs BRME1-N | undetectable |  |  |  |  | 303 |
| HSF2BP vs BRME1-C | undetectable |  |  |  |  | 293 |
| HSF2BP vs BRME1-C | undetectable |  |  |  |  | 303 |
| HSF2BP $\alpha 1$ vs BRME1-M | 2.5E-08 (4.5) | 0.96 | -7.6 (0.06) | -10.5 | -2.90 | 303 |
| HSF2BP $\alpha 1$ vs BRME1-M | 2.1E-08 (4) | 0.93 | -1.2 (0.05) | -10.2 | -9.00 | 293 |
| HSF2BP+BRME1-M vs BRCA2-HBD | 1.3E-08 (2) | 0.37 | -10.5 (0.13) | -10.4 | -0.10 | 293 |
| HSF2BP+BRME1-M vs BRCA2<sub>2213-2342</sub> | 7.8E-09 (1.8) | 0.35 | -10.9 (0.12) | -12.6 | -1.70 | 293 |
| HSF2BP+BRCA2-HBD vs BRME1-M | 3.4E-08 (8) | 0.87 | -4.4 (0.06) | -10 | -5.60 | 293 |
| HSF2BP+BRCA2-HBD vs BRME1-M | 2.0E-08 (19) | 0.8 | -5.8 (0.22) | -10.2 | -4.40 | 293 |

To elucidate how HSF2BP helix α1 interacts with BRME1-M, we crystallized HSF2BP helix α1 both alone and bound to BRME1-M. We solved the crystal structure of the human HSF2BP fragment from E19 to G48. Each monomer consists of a β-strand (residues F20-R24) and an α-helix (residues K25-L46) (Extended Data Fig. 5b), as predicted by AlphaFold (Extended Data Fig. 3a,c). The crystal asymmetric unit contains two peptides interacting through antiparallel β-strands and parallel α-helices that assemble into a tetramer by application of a 2-fold crystallography symmetry (Supplementary Table 1). This tetrameric structure is similar to the best model proposed by AlphaFold, which does not fit into the cryo-EM map (Extended Data Fig. 3c). We also solved the crystal structure of helix α1 bound to BRME1-M. Each BRME1-M monomer contains a large α-helix (residues T605-R638) (Figure 4d). The asymmetric unit contains a HSF2BP helix α1 and a BRME1-M peptide that forms a parallel heterodimer. Two of these dimers assemble into a parallel tetramer by application of a 2-fold crystallographic symmetry (Supplementary Table 1). The β-strands from the two HSF2BP peptides form an anti-parallel β-sheet. Altogether, our X-ray crystallography analyses provided an antiparallel structure for the α1 tetramer and a parallel arrangement for the helices of the α1-BRME1 heterotetramer. Our cryo-EM data and AlphaFold modelling suggested that homotetramerization of α1 in a parallel configuration is responsible for the formation of the ring. Based on the X-ray structure of α1 bound to BRME1-M in addition to a previous study^16^, we further hypothesized that BRME1-M α-helix competes with HSF2BP helix α1, thus disassembling α1 tetramers. To experimentally validate this hypothesis, we performed mass photometry analyses on HSF2BP in the absence and presence of BRME1-M. We observed that indeed, the HSF2BP tetramer is dissociated into dimers after addition of BRME1-M (Fig. 4e).

### BRME1 binding to HSF2BP dissociates HSF2BP-BRCA2 ring complexes

We then proceeded to determine the effect of BRME1 on the structure of the HSF2BP-BRCA2 complex. We observed using negative-staining EM that addition of BRME1-M to a 2:1 mix of HSF2BP and BRCA2-HBD resulted in a near-complete disappearance of the large ring-shape complexes (Fig. 5a). To replicate this dramatic effect under different experimental conditions and to quantify it, we performed single-molecule analysis using scanning force microscopy (SFM; Fig. 5b-d). HSF2BP alone appeared on SFM scans as round objects with a volume distribution centered around 95 and 190 nm^3^. We surmised that under the low protein concentrations used for single-molecule SFM experiments, HSF2BP oligomer equilibrium may be shifted towards dimers and interpreted the two populations of objects as tetramers that, respectively, dissociated during deposition or remained intact (Fig. 5b,c; Extended Data Fig. 6a,b). This interpretation is consistent with the mass photometry data recorded on HSF2BP at different concentrations, which revealed that, whereas at 50 nM HSF2BP is a tetramer (Fig. 4e), at 25 nM it is a mix of dimer and tetramer (Extended Data Fig. 6c). We also observed by SFM that the armadillo domain alone appeared as smaller round objects with a volume distribution centered on 35 and 90 nm^3^, that we assigned to monomers and dimers, respectively (Fig. 5d). By contrast, the volume distribution of the complexes formed by HSF2BP and BRCA2-HBD revealed significantly larger objects, with peaks at ∼550 and ∼2,000 nm^3^ (Fig. 5b,c; Extended Data Fig. 6a,b). Taking the 95 nm^3^ volume of HSF2BP dimers as the smallest unit and correcting for the increase in measured volume due to the ringed shape and scanning precision, we assigned the minor sub-population of the largest objects (∼2,000 nm^3^) to the 24-mers. This was confirmed by analyzing the sample of HSF2BP-BRCA2-HBD that was purified by Size-Exclusion Chromatography (SEC) and used in the EM analyses (Fig. 5d). Addition of BRME1-M to the HSF2BP-BRCA2-HBD sample used for the first SFM measurements resulted in a significant reduction in apparent object size, which, when quantified, corresponded to a complete disappearance of the 24-mer population (Fig. 5b,c; Extended Data Fig. 6a,b). Octameric species were still observed, and the number of tetramer- and dimer-sized objects was increased.

**Fig. 5.**
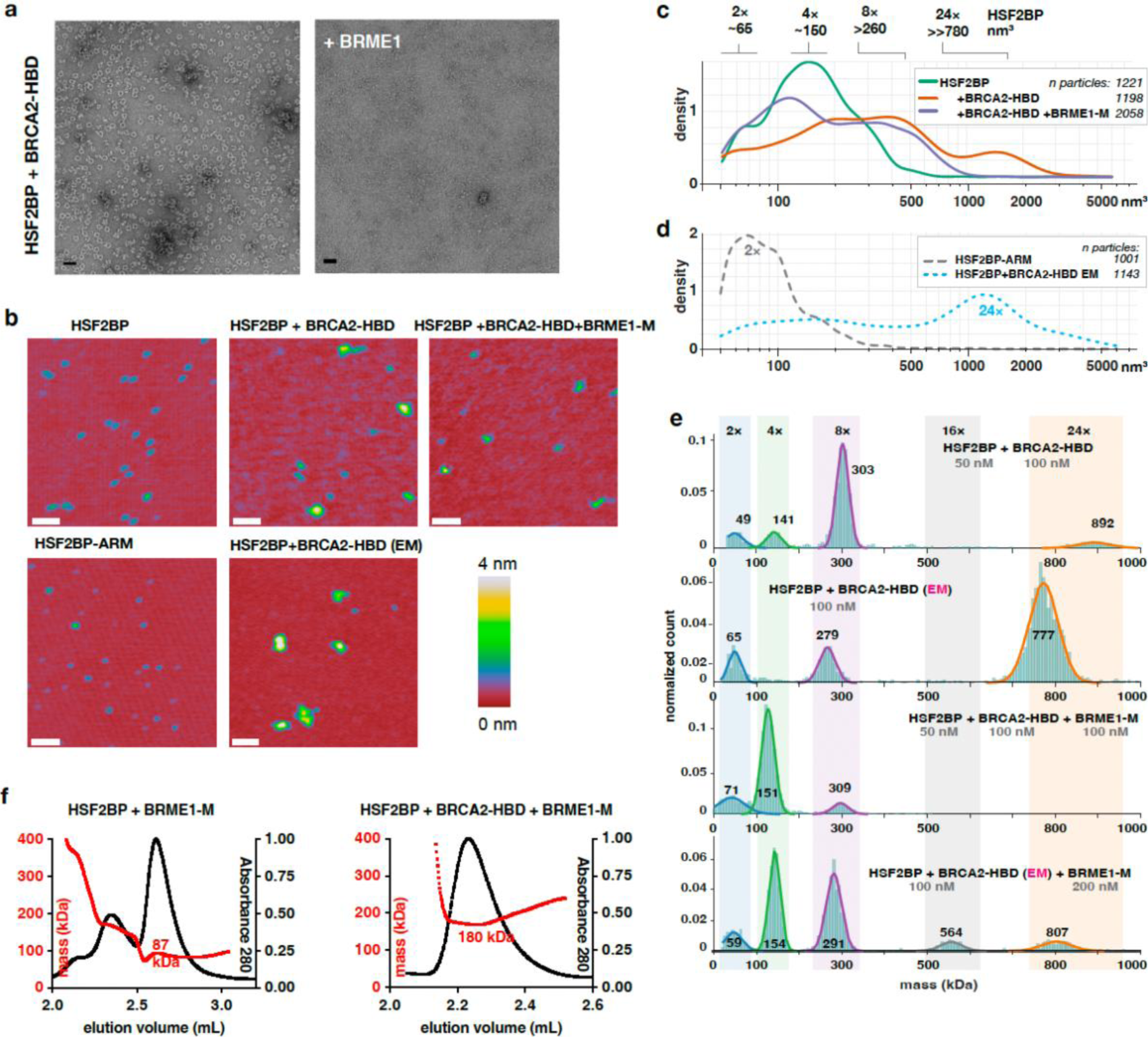
A BRME1 peptide disrupts the ring-shaped complex. **a**, Negative staining EM images recorded on the complex formed by HSF2BP and BRCA2-HBD in the absence (left) and presence (right) of the peptide BRME1-M. The conditions are the same as in Fig. 2c. **b**, Representative SFM images of HSF2BP, its armadillo domain (HSF2BP-ARM), HSF2BP+BRCA2-HBD and HSF2BP+BRCA2-HBD+BRME1-M. The complexes were assembled just before deposition on mica for SMF analysis. Only the sample prepared for EM was purified by gel filtration. Scale bar = 100 nm. **c**, Kernel density plots of the particle volume distributions for HSF2BP and its complexes with BRCA2-HBD and BRCA2-HBD+BRME1-M. The number of particles is indicated in the legend, the replicate of the experiment is shown in Extended Data Fig. 6a,b. **d**, Kernel density plots of particle volume distributions of the HSF2BP-ARM fragments and HSF2BP+BRCA2-HBD complexes prepared following the protocol used for EM. These were used to determine the volumes of the dimers and the ring-shaped 24-mer, respectively. The experiment was done once, number of analyzed particles is indicated. **e**, Mass photometry experiments performed on HSF2BP in the presence of BRCA2-HBD (freshly assembled or cryo-EM samples) and/or BRME1-M. The distributions of masses are displayed at the indicated concentrations of proteins. The cryo-EM sample was diluted down to 100 nM and a 2-fold excess of BRME1-M was used to observe the impact of BRME1-M on the 24mer complex. Replicates are presented in Extended Data Fig. 6d. **f**, SEC-MALS analysis of the complex between HSF2BP and BRME1-M, in the absence (left) or presence (right) of BRCA2-HBD. Replicates are presented in Extended Data Fig. 6e.

To support these SFM results, we performed additional mass photometry analyses of both freshly prepared samples of BRCA2-HBD bound HSF2BP and the sample used for cryo-EM studies. We observed that, at the concentrations used, addition of BRCA2-HBD to HSF2BP triggers the formation of octamers and, in smaller amounts, 24-mers (Fig. 5e; Extended Data Fig. 6d). Further addition of BRME1-M disassembles the 24-mer complexes, yielding to a mix between octamers, tetramers and dimers. Similarly, addition of BRME1-M to the cryo-EM sample caused disassembly of the ring-shaped complex, and enrichment in double-octamers, octamers and tetramers (Fig. 5e). Altogether, our EM, SFM and mass photometry results consistently demonstrated that BRME1 can act as a dissociation factor for HSF2BP-BRCA2 oligomers.

Finally, we characterized the complex between HSF2BP, BRCA2-HBD and BRME1-M using SEC-MALS. We first purified HSF2BP bound to BRME1-M and verified using SEC-MALS that it is a dimer (Fig. 5f; Extended Data Fig. 6e). We then prepared a sample of the ternary complex at the highest possible concentration (1 mg/ml, the solubility limit of this complex in our conditions) in the presence of an excess of BRCA2-HBD and BRME1-M peptides, and performed a SEC-MALS analysis on this sample. We observed that the ternary complex has a molecular mass of 181 ± 1 kDa (n=2) (Fig. 5f; Extended Data Fig. 6e). The theoretical mass of four HSF2BP bound to four BRME1-M and two BRCA2-HBD being 182 kDa, we concluded that this complex contains a tetrameric form of HSF2BP.

### BRME1 protects cancer cells from BRCA2 inhibition by HSF2BP

In our previous work, we demonstrated that ectopic HSF2BP compromises BRCA2 function and causes BRCA2 proteasomal degradation. This degradation involved the p97 segregase ^24^, an hexameric protein that unfolds and disassembles ubiquitylated substrates, to pull proteins out of membranes, segregate proteins from partners for downstream activity or unfold proteins for proteasomal degradation ^28^. Given the dissociative effect of BRME1 on HSF2BP-BRCA2 multimers, we hypothesized that de-aggregation by BRME1 will cancel the effect of HSF2BP. To test this hypothesis, we stably produced in HeLa cells HSF2BP alone and together with BRME1 (Fig. 6a,b) and measured their effect on resistance to DNA inter-strand crosslinking agents and PARP inhibitors (Fig. 6c-e). BRME1 on its own had no effect on the cell sensitivity to DNA damage, but it completely abolished the sensitization induced by HSF2BP, consistent with our hypothesis. The C-terminal HSF2BP-binding part of BRME1 had the same protective effect as the full-length protein, demonstrating that disrupting BRCA2-HSF2BP multimerization may be sufficient to protect BRCA2 function. We also found that the HSF2BP variant lacking the tetramerization-mediating helix α1 does not sensitize cells. Taken together, these data suggested that HSF2BP-BRCA2 oligomerization compromises BRCA2 function and that BRME1 prevents this by dissociating the oligomers.

**Fig. 6.**
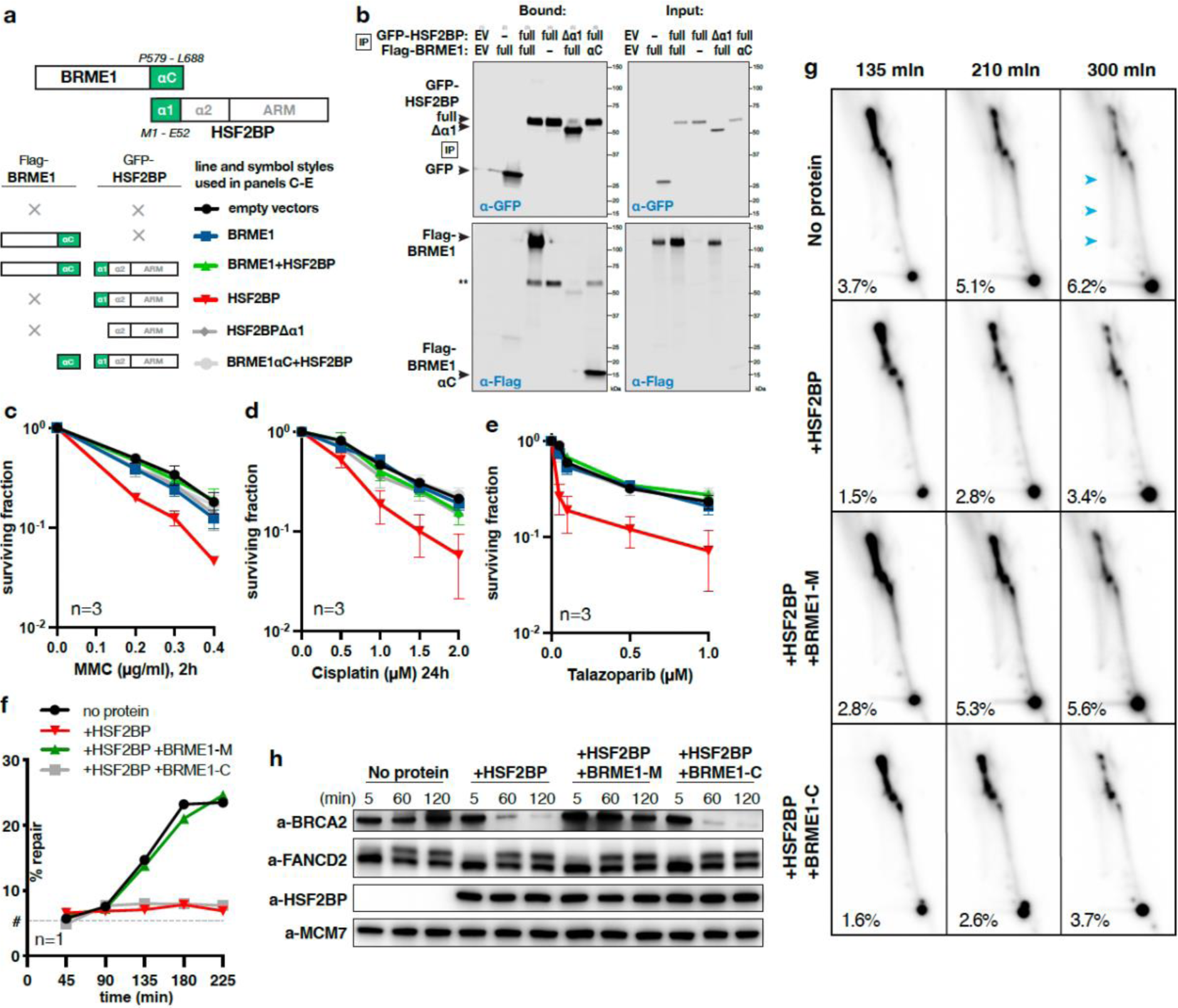
BRME1 protects cancer cells from HSF2BP and prevents BRCA2 degradation during interstrand crosslink repair. **a**, Schematic depiction of the HSF2BP and BRME1 fragments used in the congenic survivals shown in panels **c-e.** Interacting α-helices are shown as green blocks, the six combinations of HSF2BP and BRME1 variants used in the survivals are shown next to the corresponding line styles used in panels (**c-e**); × indicates that the protein was not present. **b**, Co-immunoprecipitation of HSF2BP and BRME1 variants used in clonogenic survivals. Proteins eluted from the anti-GFP beads were detected by immunoblotting with the indicated antibodies. **c-e**, Clonogenic survival of HeLa cells stably producing HSF2BP and BRME1 variants, as indicated in panel **a**. Cells were treated with mitomycin C (MMC), cisplatin or talazoparib. The experiments were repeated three times, means and s.e.m. are plotted. **f**, Efficiency of synthetic cisplatin interstrand crosslink repair in *Xenopus* egg extract in the presence or absence of HSF2BP, the BRME1 peptides BRME1-M (binding HSF2BP) and BRME1-C (not binding HSF2BP; Fig. 4a,b). See Sato *et al.* 2020 for the detailed description of the assay ^24^. Replicates of the experiment are shown in Extended Data Fig. 7c,d. **g**, HR intermediate formation during the repair of synthetic cisplatin DNA interstrand crosslink in Xenopus egg extract monitored by 2D agarose gel electrophoresis. The reactions were carried out in the presence or absence of HSF2BP, the BRME1 peptides BRME1-M and BRME1-C (see Fig. 4a,b). The X-arc that contains HR intermediates is indicated by blue arrows in the upper right panel, percentage of the signal localizing to it is indicated, as detailed previously ^24^. A replicate of this experiment is shown in Extended Data Fig. 7e. **h**, Effect of HSF2BP and BRME1-M or BRME1-C peptides on the endogenous *Xenopus* BRCA2 protein during the time course (5-120 min) of the interstrand crosslink repair reaction. Antibodies used for immunoblotting are indicated. See Sato *et al.* 2020 for the validation and analysis of the dependencies of BRCA2 protein degradation induced by HSF2BP ^24^. A replicate of this experiment is shown in Extended Data Fig. 7f.

To further distinguish the function of octamers versus 24-mers, we tested the HSF2BP variant H80A-R84A that exclusively forms octamers when bound to BRCA2-HBD (Fig. 3d). Measuring the impact of overexpressing this variant in HeLa cells showed that, upon DNA damage, HSF2BP H80A-R84A compromised cell survival as HSF2BP wild-type (Extended Data Fig. 7a,b). These results suggested that the HSF2BP capacity to induce BRCA2 unloading from the DNA damage site and degradation necessitates the formation of HSF2BP octamers, but not 24-mers.

### BRME1 prevents HSF2BP-induced BRCA2 degradation

To establish the mechanism by which HSF2BP attenuates BRCA2 function, we previously studied the repair of a single chemically defined and site-specific cisplatin DNA inter-strand crosslink in a plasmid replicating in *Xenopus* egg extract ^24^. These biochemical experiments showed that both *Xenopus* and human HSF2BP inhibits replication-dependent restoration of the genetic information at the crosslink as measured by the regeneration of a restriction enzyme site. We demonstrated using two-dimensional agarose gel electrophoresis that the HR step and not the preceding steps (recognition, signaling, crosslink unhooking by nucleases, or translesion synthesis) of the reaction was inhibited, and that the immediate reason for the inhibition was HSF2BP-induced BRCA2 degradation. To determine whether the protective effect of BRME1 we observed in cancer cells ectopically producing HSF2BP would also manifest biochemically, we performed the same *Xenopus* egg extract assays (Fig. 6f-h; Extended Data Fig. 7c-f) in the presence of either the BRME1-M peptide, which binds HSF2BP α1, or the adjacent BRME1-C peptide, which does not bind HSF2BP (Fig. 4a,b). Consistent with the cancer cell data, BRME1-M reverted the inhibitory effect of HSF2BP on the interstrand crosslink repair reaction (Fig. 6f), and specifically suppressed the reduction in the formation of HR repair intermediates monitored by two-dimensional agarose DNA gel electrophoresis (Fig. 6g); the non-binding BRME1-C peptide had no such effect. Moreover, HSF2BP-induced BRCA2 degradation was completely abolished by the BRME1-M but not the BRME1-C peptide (Fig. 6h). Combined with the structural and cancer cell data, this suggests that BRME1 acts as a disaggregation factor for HSF2BP-BRCA2 complexes, preventing their recognition by the p97/VCP segregase and degradation by the proteasome.

## DISCUSSION

The HR mediator BRCA2, as well as the newly described meiotic proteins HSF2BP and BRME1, participate to meiotic HR. They are essential for male fertility. Recent studies revealed that, in germline cells, BRCA2 functions in a tight, likely constitutive, complex with HSF2BP and BRME1. BRCA2 directly binds to HSF2BP via a unique repeat-mediated oligomerization-inducing mechanism. BRME1 also directly binds to HSF2BP and phenocopies it. Here, we extended our structural characterization of this germline BRCA2 complex ^23^, by studying the full-length HSF2BP, rather than only its armadillo domain, in complex with several BRCA2 and BRME1 peptides. Our findings support our previous proposal that HSF2BP can act as a polymerization factor for BRCA2, thanks to three oligomerization mechanisms: two intrinsic to HSF2BP (homo-oligomerization via α1 and dimerization via α2+armadillo) and one extrinsic, mediated by the repeats encoded by exons 12 and 13 of BRCA2 (Fig. 7). In addition, our study reveals several properties that we did not anticipate from the previous analyses, most importantly: tetramerization of HSF2BP into a V-shaped structure revealed by SAXS, and formation of a large ring-shaped 880 kDa hetero-oligomer consisting of 24 HSF2BP and 12 BRCA2-HBD molecules organized into three interlocked diamond-shaped octameric (8×HSF2BP + 4×BRCA2-HBD) rings, revealed by cryo-EM and supported by SFM, SEC-MALS, AlphaFold modelling and X-ray crystallography (Fig. 1-3). We also delineated a BRME1 α-helical peptide that can bind and displace helix α1 of HSF2BP, thus disrupting the tetramer formed by helix α1 in HSF2BP and the 24-mer complex formed by full-length HSF2BP upon binding to BRCA2-HBD (Fig. 4, 5). Altogether, our results provide a molecular mechanism for the aggregator-disaggregator model explaining the apparently opposite effects of HSF2BP on BRCA2-mediated HR in somatic vs germline cells.

**Fig. 7.**
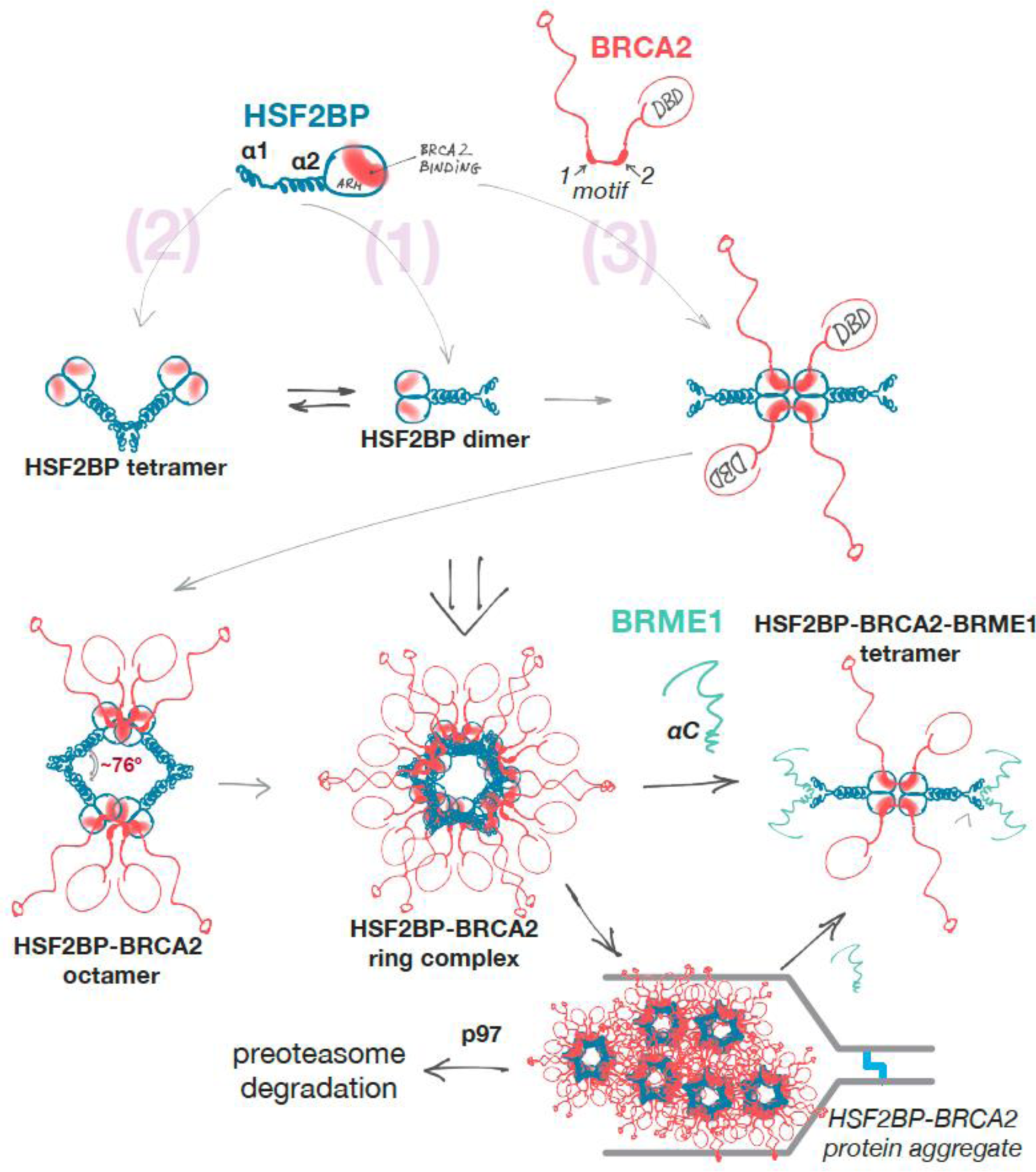
Oligomeric states of HSF2BP and the “aggregator-disaggregator” model proposed to explain the effects of HSF2BP and BRME1 on BRCA2 in somatic and germline cells. Oligomers observed experimentally are indicated in bold. Two intrinsic (1) (2) and one BRCA2-mediated (3) oligomerization mechanisms result in the formation of a constitutive homodimer, a V-shaped tetramer and a large ring complex. This concentrates and organizes BRCA2, but at replication fork stalled at crosslink, it results in aggregation and proteasomal degradation of BRCA2. BRME1 competes with one of the oligomerization mechanisms (2), resulting in the formation of a 4×HSF2BP:2×BRCA2:4×BRME1 complex and thus preventing or reversing BRCA2 aggregation.

The remarkable ring structure of the HSF2BP-BRCA2 complex revealed by our cryo-EM analysis comes from two new properties of HSF2BP we identified: (1) HSF2BP is a tetrameric protein with a V-shape, the angle at the bottom of the V being defined by the α1-mediated tetramerization constraint; (2) in the ring-shaped HSF2BP-BRCA2 complex, the α2 coiled coils and armadillo domains of the three octamers interact through interfaces that are at least partially evolutionary conserved (Extended Data Fig. 8a,b). A relatively constrained V-shape favors the formation of an HSF2BP octamer upon binding to the BRCA2 peptide. In this octamer, the α2 coiled coils lie in the same plane. They outline a diamond shape, the angle at the vertex formed by α1 helices (∼76°) being favored by the HSF2BP tetramerization mode revealed in this study and the supplementary angle at the vertex formed by the armadillo domains (∼104°) being defined by the BRCA2-dependent tetrameric structure described previously ^23^. The structural origin of the angle value at the α1 vertex is not clear and it may have a degree of flexibility in solution, but we postulate that such angle value must be constrained to explain why HSF2BP bound to BRCA2-HBD organizes into closed diamond-shaped octamers rather than linear polymers. These octamers could not be structurally characterized independently, as three of them interlock to form the observed ring, which is the preferred conformation for the complex at high concentration (μM). Evolutionary conservation of the complex, which allowed us previously to use human and frog HSF2BPs interchangeably in biochemical experiments, suggested an explanation to this. Contacts between the interlocked octamers are mainly mediated by helices α2: in each octamer, two α2 coiled coils interact with a second octamer, whereas the two other α2 coiled coils interact with the third octamer. While the details of these interfaces are missing, we have identified a double mutant, H80A-R84A, which only forms octamers upon binding to BRCA2-HBD (Fig. 3d). H80 and R84 are conserved through evolution (Extended Data Fig. 9a). They are not essential for the disruptive function of HSF2BP in cancer cells. However, their role might be important in meiosis. The other conserved interfaces of the ring-shaped complex are the groove binding to BRCA2-HBD, the α1 regions involved in tetramerization and the armadillo regions involved in dimerization (Extended Data Fig. 8a). We also noted the presence of three cysteine pairs located at the HSF2BP dimerization interface and were curious whether they are required for HSF2BP function and could be a point of redox regulation (Extended Data Fig. 9a-d). The triple C78A-C120S-C128A substitution variant sensitized HeLa cells less than the wild-type protein. The C128R variant was also found in a family with premature ovarian insufficiency^29^. Altogether, we surmise that (i) conservation of the interfaces reflects the functional importance of the ring, (ii) dimerization of HSF2BP is essential for most of its functions, and (iii) formation of the octamers upon binding to BRCA2-HBD is important for the disruptive function of HSF2BP in cancer cells.

The distinct structure of the HSF2BP-BRCA2 ring complex makes it tempting to speculate how such spatial organization might be functionally relevant. There are multiple precedents for the specific role of ring-shaped complexes in DNA metabolism and HR specifically, from sliding clamps and hexameric helicases to the octameric ring formed by DMC1. However, the ring we characterized is larger than the ring complexes that encompass DNA: it has an inner diameter of 100 Å compared to 25 Å for hexameric helicases or 35 Å for PCNA ^30, 31^. Also, the inner surface of HSF2BP-BRCA2 is negatively charged (Extended Data Fig. 8c), and HSF2BP has no detectable affinity for DNA ^23^, whereas the inner surfaces of the DNA-encompassing rings are positively charged. Given the proposed role of HSF2BP at resected meiotic DSBs coated by ssDNA-binding proteins RPA, SPATA22 and MEIOB and the reported interactions between HSF2BP and these proteins, it is notable that the opening of the ring is sufficiently large to accommodate protein-coated ssDNA (∼40-80 Å ^32, 33^). It also notable that BRCA2-HBD molecules are all exposed on the outer surface of the ring, so that BRCA2 might partially hinder access to HSF2BP, but should allow threading of protein-DNA complexes through the opening.

The major issue for these hypotheses is the disruptive role of BRME1 on the ring structure, and the strong phenotypic and cytological similarity between BRME1 and HSF2BP mutants, which suggests that the two proteins function as a complex in meiosis. In this context, we propose that, in the presence of BRME1, an alternative HSF2BP tetrameric conformation is formed, linked by two BRCA2 molecules and blocked by four BRME1 molecules from forming higher-order assemblies. This is supported by the appearance of particles with close to tetrameric volumes, as observed in our SFM and mass photometry experiments, in the presence of both BRCA2-HBD and BRME1-M (Fig. 5c,e). It is also supported by SEC-MALS experiments performed at higher concentration and in the presence of excess of peptides, in which only a tetramer is observed (Fig. 5f). Such tetramer may carry out some or all of the known HSF2BP functions in meiosis. It has a 2-fold symmetry and could bridge two other entities as has been shown for monopolin, a V-shaped dimeric protein bridging kinetochores during yeast meiosis ^34^. However, it is possible that BRME1 serves as a regulatory factor that ensures that the HSF2BP-BRCA2 ring is assembled at the right time and place, preventing ectopic BRCA2 polymerization that can have detrimental effects, as revealed by our study of cancer cells discussed below. The release of BRME1 from HSF2BP could be controlled by post-translational modification or other mechanisms, such as competition with partners. In this regard, it is remarkable that HSF2BP α1 can both homo- and heterotetramerize. Most helix α1 surface is evolutionarily conserved (Extended Data Fig. 8a; Extended Data Fig. 9a), suggesting that both homo- and hetero-tetramerization are important for HSF2BP function. Taken together, we hypothesize that organizing BRCA2 into a ring-shaped oligomer that can encompass objects the size of protein-coated ssDNA is an important feature of HSF2BP that is under positive evolutionary selection.

While the relevance of the ring-shaped BRCA2-HSF2BP complex in meiosis is speculative at the moment, we suggest that our findings resolve the arguably most puzzling observation about HSF2BP, namely its apparently opposite effects on somatic vs meiotic HR (Fig. 6, 7). We previously found that ectopic HSF2BP in human cancer cells or *Xenopus* egg extracts suppresses HR during interstrand crosslink repair by inducing BRCA2 degradation. This degradation is dependent on the p97 segregase, which processes protein aggregates ^23^. We hypothesized that the polymerizing effect of HSF2BP on BRCA2 turns into aggregation in ectopic settings, thus inducing p97-dependent BRCA2 degradation. It was however not clear what restricts this polymerization both spatially, stopping the formation of endless HSF2BP-BRCA2 chains via α1- and BRCA2-linked armadillo connections, and temporally, restraining it to specific ectopic contexts. Circularization into octamers and ring complexes allows HSF2BP-BRCA2 to diffuse through the cytoplasm without forming linear chain aggregates. The disaggregating effect of BRME1 explains why aggregation does not happen in testis and ES cells. Aggregation followed by proteasomal degradation happens only at the stalled replication fork. It may be triggered by high local concentration of the proteins in foci, additional oligomerizing interactions in these foci with other proteins or local post-translational modifications. Alternatively, the aggregation-sensing machinery may be more active at the crosslink-blocked fork, which is known to recruit ubiquitin ligases and the VCP/p97 segregase itself. BRME1 was not present in the cancer cells that we used during our first analyses, explaining why we initially observed a disruptive effect for HSF2BP. Differential expression of HSF2BP and BRME1, as well as alternative splicing of *BRCA2* exon 12 ^17, 23, 35–37^, provide cancer cell with an evolutionary platform to drift between genomic instability and drug resistance.

In conclusion, HSF2BP can polymerize BRCA2 into a large ring-shaped structure, consisting of three interlocked BRCA2-bound HSF2BP octamers. Formation of this complex is regulated by BRME1. Under specific DNA damage conditions, ectopic expression of HSF2BP in cancer cells can trigger BRCA2 aggregation and degradation, resulting in genomic instability. When HSF2BP is expressed physiologically, along with BRME1, aberrant formation of this complex is prevented. We propose that the ring-shaped complex timely concentrates and organizes BRCA2 and other bound proteins and can accommodate ssDNA coated with ssDNA binding proteins, in order to facilitate BRCA2-mediated DSB repair in meiocytes.

## METHODS

### Protein expression and purification

Human HSF2BP, either full-length wild-type, its substitution variants (H80A-R84A; Q104A-E108A) or the truncation variant lacking helix α1 (from G48 to V334), was expressed using a pETM11 (6×His-TEVsite) expression vector in *E. coli* BL21 (DE3) Rosetta2 pLysS or Star strains. A starter culture (LB + 50 μg/ml kanamycin, 30 μg/ml chloramphenicol) was grown overnight at 37 °C, and used to inoculate 1 L of LB which was grown at 37 °C until the OD_600_ reached 0.6. Expression was induced by addition of 0.2 mM IPTG, and continued at 20°C overnight. Harvested cells were resuspended in 25 mM Tris-HCl pH 8, 500 mM NaCl, 5 mM β-mercaptoethanol, EDTA-free Protease Inhibitor Cocktail (Roche) and disrupted by sonication. Lysates were supplemented with 1 mM MgCl_2_, treated by benzonase nuclease at 4 °C for 30 min, and then centrifuged at 15.000 × g at 4 °C for 30 min. After filtration (0.4 µm), the supernatant was loaded on a chromatography HisTrap FF crude 5 mL column (GE Healthcare) equilibrated with Tris-HCl 25 mM pH 8, 500 mM NaCl and 5 mM β-mercaptoethanol. HSF2BP (variant) was eluted with a linear gradient of imidazole. The tag was cleaved by the TEV protease (at a ratio of 2% w/ w) during an ON dialysis at 4 °C against 25 mM Tris-HCl pH 7.5, 250 mM NaCl, 5 mM β-mercaptoethanol. The protein solution was loaded on a HisTrap column and the tag-free HSF2BP was collected in the flow through. Finally, a size exclusion chromatography was performed on HiLoad Superdex 10/300 200 pg equilibrated in 25 mM Tris pH 7.5, 250 mM NaCl, 5 mM β-mercaptoethanol. The quality of the purified protein was analyzed by SDS-PAGE and the protein concentration was determined by spectrophotometry from the absorbance at 280 nm.

The gene coding for BRCA2-HBD (BRCA2 residues 2291 to 2342, including mutation C2332T to avoid oxidation problems) was optimized for expression in bacteria and synthesized by Genscript. It was cloned in a pET-22b vector. This vector was used to express a fusion protein comprising BRCA2-HBD, a TEV site, GB1 and a 6xHis tag in BL21 DE3 Star cells. BRCA2-HBD was purified as previously reported ^23^.

Peptides coding for HSF2BP helix α1 (residues 19 to 50) and BRME1 C-terminal region (BRME1-N: residues 578 to 601; BRME1-M: 602 to 641; BRME1-C: 649 to 668) were purchased from Genecust.

The complex between HSF2BP and BRCA2-HBD was prepared by mixing the two proteins at a molar ratio of 1:1.5. The mix was concentrated by centrifugation at 4500 ×g using a 3 kDa cutoff membrane at 4°C, and then loaded on a size exclusion column Superdex 200 increase 10-300 GL (GE Healthcare) equilibrated with 25 mM Tris-HCl pH 7.5, 150 or 250 mM NaCl, 5 mM β-mercaptoethanol. The peak fractions were collected and further analyzed by EM.

The thermal stability of HSF2BP either free or bound to BRCA2-HBD was evaluated using the simplified Thermofluor assay available on the High Throughput Crystallization Laboratory (HTX Lab) of the EMBL Grenoble ^38^. Samples were analyzed at different concentrations: HSF2BP at 7 mg/ml and 14 mg/ml; HSF2BP bound to BRCA2-HBD at 1 mg/ml and 12 mg/ml.

### Mass Photometry

Mass photometry experiments were recorded on a Refeyn Two^MP^ system. Samples were prepared at 1 μM and diluted in 25 mM Tris-HCl buffer pH 7.5, 250 mM (or 150 mM when indicated) NaCl, 5 mM β-mercaptoethanol before the experiments to reach concentrations comprised between 25 and 100 nM. A two-fold excess of peptides (either BRCA2-HBD or BRME1-M) was used to measure the impact of peptide binding on HSF2BP oligomerization. Calibration of the instrument was performed in the same buffer with carbonic anhydrase, BSA, and urease covering a range of molecular weights from 29 kDa to 816 kDa. Data were analyzed using the provided Discover analysis software. Masses were estimated by multi-Gaussian fitting using the online PhotoMol software (https://spc.embl-hamburg.de/app/photoMol) ^39^.

### SEC-MALS

Size-exclusion chromatography (SEC) coupled to multi-angle light scattering (MALS) was used in order to measure the molecular masses of the complexes in solution. Therefore, HSF2BP (full-length wild-type or its substitution and truncation variants) was loaded in the presence or absence of BRCA2-HBD and BRME1-M, on a Superdex 200 10/300 GL (GE Healthcare; flow rate: 500 μl/min, data in Fig. 1b, 2a) or a BIOSEC 3 column (Agilent; flow rate: 200 μl/min, all other experiments), using a HPLC Agilent system coupled to MALS/QELS/UV/RI (Wyatt Technology). The chromatography buffer was 25 mM Tris-HCl buffer, pH 7.5, 250 mM NaCl, 5 mM β-mercaptoethanol. The proteins were injected at 0.8-7 mg/ml in 20-100 μl. Data were analyzed using the ASTRA software; a calibration was performed with BSA as a standard. To represent the data, the normalized absorbance at 280 nm was overlaid with the molar mass (Da), and both parameters were plotted as a function of the elution volume.

### SEC-SAXS

SEC coupled to small angle X-ray scattering (SAXS) is available on the SWING beamline at synchrotron SOLEIL, in order to obtain a distance distribution corresponding to each sample in solution. The free HSF2BP protein was analyzed using a BIOSEC 3 column (Agilent) equilibrated in 25 mM Tris-HCl pH 7.5, 250 mM NaCl, and 5 mM β-mercaptoethanol. The protein was loaded at a concentration of 6 mg/ml, in order to observe an elution peak at an OD_280nm_ of 1 AU. The SAXS intensity was measured every second during the elution as a function of the scattering angle. The final SAXS curve is the sum of the curves recorded on the elution peak. A distance distribution curve P(r) was calculated, whose Fourier transform fits to the SAXS curve. The residual values are the experimental minus the fitted SAXS intensity values, divided by the experimental errors. The distance distribution P(r) was plotted in arbitrary units as a function of the distance, the maximal distance being 250 Å.

### Sample preparation for EM

The quality of the EM sample was checked by negative-staining EM. Therefore, the complex between HSF2BP and BRCA2-HBD was diluted to approximately 0.05 mg/ml in 25 mM Tris-HCl pH 7.5, 250 mM NaCl. Three microliters were deposited on an airglow-discharged carbon-coated grid. Excess liquid was blotted, and the grid rinsed with 2% w/v aqueous uranyl acetate. The grids were visualized at 100 kV with a TECNAI Spirit (FEI) transmission electron microscope (ThermoFisher, New York NY, USA) equipped with a K2 Base 4k × 4k camera (Gatan, Pleasanton CA, USA). A non-diluted sample (2-4 mg/ml) concentrated in 25 mM Tris-HCl pH 7.5, 150 mM NaCl, 5 mM β-mercaptoethanol was further prepared for cryo-EM. An aliquot of 2.5 µl was deposited onto freshly glow-discharged, Quantifoil R2/2 300-mesh Gold grids. These grids were frozen in liquid ethane using a Vitrobot Mark IV (ThermoFischer) at 100 % humidity, 4 °C, using a blotting force of 1 and a blotting time of 5 s.

### Cryo-EM data acquisition and image processing

For cryo-EM analysis, 9531 movies were collected using the EPU software on a Titan Krios electron microscope (Thermo Fisher Scientific) operating at 300 kV, equipped with a Selectris X filter, slit open at 10 eV and a Falcon 4i direct electron detection camera (Supplementary Table 2). The defocus range was between −0.4 and −1.8 µm, with a pixel size of 0.73 Å. The total dose was 40 electrons/Å^2^ distributed on 40 frames. Movie frames were aligned using MotionCor2 ^40^, with applied dose compensation. Motion-corrected micrographs were imported into CryoSPARC ^41^. Contrast transfer function (CTF) parameters were estimated using CTFFIND4 ^42^.

Micrographs were examined and those with a CTF fit resolution higher than 5.5 Å or a relative ice thickness larger than 1.2 were excluded. The remaining 8962 micrographs were used for blob picking as described in Extended Data Fig. 2b. After several rounds of particle picking and classification, first three volumes were calculated and refined. These volumes corresponded to open and closed rings; particles used to build closed rings were selected for further classification and template picking. After several iterations including template picking, 2D classification, *ab initio* volume calculation and heterogeneous refinement, seven 2D classes were selected to train the Topaz tool ^43^, which was then able to pick a larger set of particles. These particles were enriched in side views of the complex. A minor family of particles showing two interlocked rings was observed and excluded. The remaining particles were used to calculate three new volumes, which corresponded again to open and closed rings. A set of particles consistent with closed rings were further used for a new Topaz training. Using this tool, a new set of particles was picked, and added to the first set of particles picked by Topaz. Duplicates were removed, three volumes were calculated and refined. One of these volumes had the shape of an open ring, the two other volumes were well-resolved rings. The 398 000 particles corresponding to these two volumes were used to calculate the final cryo-EM map of the complex (Extended Data Fig. 2c). Then, the 186 000 and 212 000 particles were separately used to calculate two new cryo-EM maps (Extended Data Fig. 2d). Whereas the GSFSC resolution of the first map is 3.3 Å, the resolutions of the two other maps are slightly lower (3.5-3.6 Å). However, the local resolution distribution of these two maps is different, with the map built from 212 000 particles showing overall a better resolution, and the map built from 186 000 particles showing better resolved inner armadillo tetramers but poorly resolved outer tetramers and coiled coils.

### Building and refinement of the cryo-EM model

The density map of the complex was of sufficient quality (between 3 and 8 Å) to fit the crystal structure of the armadillo domain of HSF2BP in complex with BRCA2-HBD (PDB codes: 7BDX) in ChimeraX. The map regions in which the armadillo domains were fitted showed a local resolution of 3-5 Å. In the peripheral region of the complex, corresponding to the N-terminal helices of HSF2BP, the local resolution map was in the range 5–8 Å. In these regions, AlphaFold guided model building. All figures were prepared using UCSF Chimera and UCSF Chimera X.

### Crystallization and structure determination

Synthetic peptides corresponding to helix α1 of HSF2BP (E19-V50) and BRME1-M (E602-K641) were dissolved in 25 mM Tris pH 7.5, 250 mM NaCl and 5 mM β-mercaptoethanol. The concentration of the helix α1 peptide was 10 mg/mL. This peptide was mixed with the BRME1-M peptide to reach a concentration of 5 mg/mL. Crystallization assays were carried out at 277 K in the High Throughput Crystallization (HTX) laboratory (EMBL Grenoble). Several crystallization conditions were identified within one day and the crystals were grown within two weeks. They were prepared for X-ray diffraction experiments using the CrystalDirect harvesting and processing robot ^44^. The best crystals diffracted at 1.47 Å (α1) and 1.98 Å (α1/BRME1-M) of resolution. They were obtained by sitting drop vapor diffusion against a reservoir containing 0.2 M LiCl, 0.1 M sodium acetate trihydrate, pH 5, and 20% PEG6000 for α1 and 0.1 M HEPES sodium salt, pH 7.5, 40% PEG200 for α1/BRME1-M. Diffraction data were collected on the *MASSIF*-1 beamline (EMBL Grenoble) of the ESRF synchrotron. Datasets were indexed, integrated and scaled using XDS^45^. The three-dimensional structures were solved by molecular replacement using the PHENIX Phaser software ^46, 47^ and an input coordinate file calculated by AlphaFold2 ^48^, were iteratively improved by manual reconstruction in COOT ^49^, and were refined using the PHENIX Refine and BUSTER (Global Phasing Limited) ^50^ software. A summary of crystallographic statistics is shown in Supplementary Table 1.

### Isothermal Titration Calorimetry (ITC) experiments

Using a VP-ITC Calorimeter (GE Healthcare), interactions between the HSF2BP protein (either full-length, or the peptide corresponding to helix α1) and the BRME1 peptides (BRME1-N, BRME1-M, BRME1-C) were characterized. The experiments were performed at two temperatures, 293 K and 303 K, and duplicated. The buffer was 25 mM Tris buffer, pH 7.5, 250 mM NaCl and 5 mM β-mercaptoethanol. 8-13 μM of HSF2BP (either full-length or helix α1) in the cell was titrated with 80-130 μM of BRME1 peptide (either BRME1-N, BRME1-M or BRME1-C) in the injection syringe. We also tested the interactions of HSFB2P bound to BRCA2-HBD with BRME1-M, and HSF2BP bound to BRME1-M with BRCA2 fragments. The experimental conditions were similar, with 10 μM of the complex in the cell, and either 200 μM of BRME1-M, or 55-60 μM of BRCA2 fragments in the injection syringe. In all cases, 10 µL of the syringe volume were injected every 210 s, except for the first injection which was of 2 µL and which was ignored in the final data analysis. This analysis was performed using the Origin 7.0 software provided by the manufacturer, in order to obtain the stoichiometries, equilibrium constants and thermodynamic parameters of the binding reactions. A summary of the ITC data is shown in Table 1.

### Scanning Force Microscopy (SFM)

Images were obtained on a NanoScope IV SFM (Digital Instruments; Santa Barbara, CA) operating in tapping mode in air with a type J scanner using silicon probes, ACT-W, with a tip radius <10 nm and a resonance frequency range of 200–400 kHz (AppNano, Santa Clara, CA). Images were processed using Nanoscope analysis (Bruker) for background flattening. The protein volumetric analyses were done using IMAGE SXM 1.89 (National Institutes of Health IMAGE version modified by Steve Barrett, Surface Science Research Centre, Univ. of Liverpool, Liverpool, U.K.). Kernel density analysis and plotting was done using the R software.

### Sample preparation for SFM

All dilutions, reactions and depositions were done in a buffer containing 25 mM Tris-HCl (pH 7.5), 250 mM NaCl and 5 mM β-mercaptoethanol. HSF2BP was diluted 10000 times to final concentration of 12 nM and deposited onto a freshly cleaved mica (Muscovite mica, V5 quality, EMS). After 30 sec the mica was washed with H_2_O and dried with a stream of filtered air. HSF2BP and BRCA-HBD_2288-2337_ complex was prepared by mixing 80 uM HSF2BP with 120 uM BRCA-HBD_2288-2337_, followed by incubation on ice for 15 min. One half of the reaction was diluted 10000 times, deposited onto a freshly cleaved mica, washed with H_2_O and dried with a stream of filtered air. Another half of the reaction was further incubated with BRME1 peptide (molar ratio HSF2BP:BRME1-M=1:1) on ice for 10 min. Sample was diluted 10000 times, deposited on mica, washed with H_2_O and dried with a stream of filtered air.

### Cell culture, constructs and stable cell line generation

HeLa (human cervical adenocarcinoma, female origin) cells were cultured in DMEM supplemented with 10% FCS, 200 U/ml penicillin, 200 µg/ml streptomycin. Expression constructs for stable production of GFP-HSF2BP, Flag-BRME1 and their truncation variants were engineered as described before ^17^ in the PiggyBac vectors by Gibson assembly and verified by sequencing. The constructs were co-transfected in combinations shown in Fig. 3A and together with the PiggyBac transposase expression plasmid (hyPBase ^51^) using Lipofectamine 3000 (Thermo Fisher) in 6-well plates seeded with 400,000 HeLa cells the day before. Selection with 1.5 µg/ml puromycin for GFP-HSF2BP constructs and 800 µg/ml G418 for Flag-BRME1 constructs was started two days after transfection and maintained for 8 days. The resulting resistant mixed cell population was used for clonogenic survival assays.

### Clonogenic survival assays

Clonogenic survival assays were performed in 6-well plates in technical duplicates. Untreated control wells were seeded with 100 cells per well in 2 mL media. Higher seeding densities of 400 and 1000 cells per well were used in the wells treated with higher drug concentrations. One day after seeding, the drugs were added: mitomycin C (MMC, Sigma, M4287-2MG), cisplatin, talazoparib (BMN-673, Axon medchem, #2502). After 2-hour (MMC) or overnight (cisplatin, talazoparib) incubation, drug-containing media was removed, wells were rinsed with PBS and refilled with 2 mL fresh media. Colonies were stained (0.25% Coomassie briliant blue, 40% methanol, 10% acetic acid) on day 10 after seeding. Plates were photographed using a digital camera, images were analyzed using OpenCFU software to quantify the colonies.

### Co-immunoprecipitation and immunoblotting

Cells were grown in 10 cm dish to near-confluence, washed twice with PBS and lysed in situ in 1 ml NETT buffer (100 mM NaCl, 50 mM Tris pH 7.5, 5 mM EDTA pH 8.0, 0.5% Triton-X100) supplemented immediately before use with protease inhibitor cocktail (Roche) and 0.4 mg/ml Pefabloc (Roche) (NETT++). After 5-10 min, cells were scraped off and collected in 1.5 ml microcentrifuge tubes; lysis was continued for additional 15-20 min on ice, then mixtures were centrifuged (15 min, 4 °C, 14000 rcf), 70 µl of the supernatant was collected as input sample, mixed with 2× Laemmli sample buffer and denatured for 5 minutes at 95 °C. The rest of the supernatant was mixed with 8 µL anti-GFP beads (Chromotek, gta-20). The mixture was incubated for 2 h at 4 °C while rotating, washed three times in NETT++ buffer and bound proteins were eluted with 70 µL 2× sample buffer, 5 min incubation at 95 °C. 15 µL bound and 7 µl input sample was separated on 4-15% TGX SDS-PAGE gel (BioRad #456-1086) and transferred to PVDF membrane (Immobilon-FL, Millipore, IPFL00010). Gels were stained with colloidal Coomassie brilliant blue (CBB) after transfer and scanned to ascertain equal loading. Immunoblotting was performed following standard procedures with anti-GFP rabbit pAb (Invitrogen, A11122) and anti-Flag mouse mAb (M2 antibody, Sigma, F3165) antibodies, followed by detection with fluorescently labeled secondary antibodies were used: anti-mouse CF680 (Sigma #SAB460199), anti-rabbit CF770 (Sigma #SAB460215). Membranes were scanned using Odyssey CLx imaging system (LI-COR).

### ICL repair assay

ICL repair assays were performed as described ^52, 53^. *Xenopus* egg extracts (HSS and NPE) and a plasmid containing a site-specific cisplatin ICL (pICL) were prepared as described previously ^54, 55^. pICL (9 ng/µl) and pQuant (0.45 ng/µl) were first incubated in a high-speed supernatant (HSS) of egg cytoplasm for 20 min at room temperature, which promotes the assembly of prereplication complexes on the DNA. Addition of two volumes nucleoplasmic egg extract (NPE) supplemented with ^32^P-α-dCTP, triggers a single round of DNA replication. Where indicated, His-tagged human HSF2BP (0.45 µM), BRME1-M peptide (1.35 µM), or BRME1-C peptide (1.35 µM) was added to NPE prior to mixing with HSS. Aliquots of replication reaction (4 µl) were stopped at various times with 45 µl Stop solution II (50 mM Tris pH 7.5, 0.5% SDS, and 10 mM EDTA,). Samples were incubated with RNase (0.13 µg/µl) for 30 min at 37 °C followed by proteinase K (0.5 µg/µl) overnight at room temperature. DNA was extracted using phenol/chloroform, ethanol-precipitated in the presence of glycogen (20 µg) and resuspended in 4 µl TE (10 mM Tris pH 7.5 and 1 mM EDTA). ICL repair was analyzed by digesting 1 µl extracted DNA with HincII, or HincII and SapI, separation on a 0.8% agarose gel in 1x TBE buffer, and quantification using Typhoon TRIO+ (GE Healthcare) and ImageQuant TL software (GE Healthcare). Repair efficiency was calculated as described ^56^.

### Two-dimensional gel electrophoresis (2DGE)

2DGE was performed as described previously ^57^. Replication intermediates of pICL at various times were extracted and digested with HincII. Fragments were then separated on a 0.4% agarose gel in 0.5× TBE buffer at 0.86 V/cm for 24 h at room temperature. The lanes of interest were cut out, cast across the top of the second-dimension gel consisting of 1% agarose with 0.3 μg/ml ethidium bromide, and run in 0.5x TBE containing 0.3 μg/ml ethidium bromide with buffer circulation at 3.5 V/cm for 14.5 h at room temperature. The gel was dried on Amersham Hybond-XL membrane and exposed to a phosphor screen. DNA was visualized using a Typhoon TRIO+.

## DATA AVAILABILITY

The coordinates and structure factors file corresponding to the crystal structures of HSF2BP helix 〈1 and the complex between HSF2BP helix 〈1 and BRME1-M described in the study were deposited in the Protein Data Bank (wwPDB) under the entry codes 8A50 and 8A51, respectively. The cryo-EM map generated in this study was deposited in the wwPDB database under accession code EMD-16432.

## ACKNOWLEDGEMENTS

We warmly thank Christophe Velours (I2BC) and Athena Collange (Synchrotron SOLEIL) for the SEC-MALS experiments; Aurélien Thureau (Synchrotron SOLEIL) for discussions about the SAXS data analysis; Florine Dupeux (IBS) and all the HTX Lab staff for the crystallogenesis and crystallography experiments; Marie-Hélène Le Du (I2BC) for helpful advice in crystallogenesis; Matthijn Vos and the NanoImaging Core at Institut Pasteur (Paris) for their support with sample preparation and image acquisition; Stéphane Bressanelli (I2BC) for his support during the cryo-EM data analyses; Bertrand Raynal and the molecular biophysics platform at Institut Pasteur (Paris) for their help during the mass photometry analyses; the staff of the I2BC computing facility (I2BC/SICS) for facilitating data management and accessibility. We acknowledge SOLEIL (SWING beamline) and ESRF (MASSIF-1 beamline) for provision of synchrotron radiation and beamlines staff for assistance in collecting data.

The research leading to these results has received funding from the European Community’s Seventh Framework Program H2020 under the project iNEXT discovery (grant agreement N°871037). It has also benefited from the cryo-EM platforms of Institut Pasteur supported by the French Government’s Investissement d’Avenir program EQUIPEX Centre d’analyse de systèmes complexes dans les environnements complexes (CACSICE, ANR-11-EQPX-008) and of I2BC supported by the French Infrastructure for Integrated Structural Biology (FRISBI, ANR-10-INSB-05) and by the CEA. This study was supported by the Oncode Institute, which is partly financed by the Dutch Cancer Society (KWF). We thank the Josephine Nefkens Cancer Program for infrastructure support. P.K. was supported by the European Research Council (ERC) through an ERC Consolidator Grant (ERCCOG 101003210-XlinkRepair) and by the Oncode Institute, which is partly financed by the Dutch Cancer Society (KWF). K.S. was supported by the Kanae Foundation for the Promotion of Medical Science and the Japanese Biochemical Society.

## AUTHOR CONTRIBUTIONS

Conceptualization, S.M., R.K., A.Z., S.Z.J.; methodology, R.G., S.M., K.S., M.O., P.L., V.R., A.A.A, A.Z.; formal analysis, R.G., S.M., K.S., P.L., C.W., P.K., A.Z., S.Z.J.; investigation, R.G., S.M., K.S., P.L., M.O., A.A.A., J.M.W., G.D., A.Z.; writing—original draft preparation, A.Z., S.Z.J.; writing—review and editing, R.G., S.M., A.Z., S.Z.J.; visualization, A.Z., S.Z.J.; supervision, S.M., C.W., P.K., R.K., A.Z., S.Z.J.; project administration, A.Z., S.Z.J.; funding acquisition, R.K., S.Z.J. All authors have read and agreed to the published version of the manuscript.

## COMPETING INTERESTS

The authors declare no competing interests.

## Notes

### Competing Interest Statement

The authors have declared no competing interest.

