## Supplementary Tables for "BRCA2-HSF2BP Oligomeric Ring Disassembly by BRME1 Promotes Homologous Recombination"

***Supplementary Table 1. Data collection and refinement statistics related to the crystal structures of HSF2BP helix $\alpha 1$ and HSF2BP helix $\alpha$1 bound to BRME1-M.***

| | HSF2BP-$\alpha$1 | HSF2BP-$\alpha$1 / BRME1-M |
|---|---|---|
| **Data collection** | | |
| Space group | $P\ 4_3\ 3\ 2$ | $I\ 4_1\ 2\ 2$ |
| Cell dimensions | | |
| *a*, *b*, *c* (Å) | 79.81, 79.81, 79.81 | 73.87, 73.87, 92.80 |
| $\alpha$, $\beta$, $\gamma$ (°) | 90, 90, 90 | 90, 90, 90 |
| *Za* | 2 | 1:1 |
| Wavelength (Å) | 0.96546 | 0.96546 |
| Resolution (Å) | 60.0 - 1.48 (1.52 - 1.48) | 36.9 - 1.9 (1.95 - 1.9) |
| Estimated resolution limit (Å)* | 1.48, 1.48, 1.48 | 2.44, 2.44, 1.71 |
| $R_{pim}$ | 0.021 (0.565) | 0.019 (0.976) |
| $R_{merge}$ | 0.050 (1.393) | 0.049 (2.384) |
| $I / \sigma I$ | 15.0 (1.2) | 12.8 (0.7) |
| $CC_{1/2}$ | 0.999 (0.559) | 0.999 (0.429) |
| Completeness (%) | 99.8 (99.9) | 99.6 (99.6) |
| Redundancy | 6.6 (6.8) | 6.9 (6.9) |
| $R_{merge}$* | 0.049 (1.049) | 0.041 (0.691) |
| $I / \sigma I$* | 15.8 (1.6) | 20.3 (3.4) |
| $CC_{1/2}$* | 0.999 (0.610) | 0.999 (0.820) |
| Completeness (%)* | 94.9 (50.6) | 61.5 (15.6) |
| **Refinement** | | |
| Resolution (Å) | 56.43 - 1.48 (1.52 - 1.48) | 36.9 - 1.9 (2.04 - 1.9) |
| No. reflections | 14159 (430) | 6522 (384) |
| $R_{work}$ / $R_{free}$ | 21.27/21.66 | 23.34/24.36 |
| No. non-hydrogen atoms | | |
| Protein | 506 | 551 |
| Ligand/ion | 10 | 38 |
| Water | 56 | 28 |
| *B*-factors | | |
| Protein | 35.8 | 52.0 |
| Ligand/ion | 21.2 | 86.0 |
| Water | 48.0 | 71.6 |
| R.M.S. deviations | | |
| Bond lengths (Å) | 0.008 | 0.008 |
| Bond angles (°) | 0.94 | 0.88 |
| PDBID | **8A50** | **8A51** |

Values in parentheses are for highest-resolution shell.
Dataset from one single crystal used per structure.
*Values calculated after truncation by STARANISO. Estimated resolution limits along the three crystallographic directions *a**, *b**, *c**.

***Supplementary table 2. Cryo-EM data collection, refinement and validation statistics.***

| | **EMD-16432** |
|---|---|
| **Data collection and processing** | |
| Voltage (kV) | 300 |
| Electron exposure (e–/Å$^2$) | 40 |
| Defocus range (μm) | -0.4 to -1.8 |
| Pixel size (Å) | 0.73 |
| Symmetry imposed | D3 |
| Initial particle images (no.) | 660 000 |
| Final particle images (no.) | 398 000 |
| Map resolution (Å) | 3.26 |
| FSC threshold | 0.143 |
| Map resolution range (Å) | 3-8 |
