## Extended data 1,2a-b for "BRCA2-HSF2BP Oligomeric Ring Disassembly by BRME1 Promotes Homologous Recombination"

**Extended Data Fig. 1. Extended data for the SEC-MALS and SEC-SAXS analyses shown in Fig. 1. a,** SEC-MALS analyses performed on HSF2BP either full-length or deleted from helix $\alpha 1$ (HSF2BP fragment from G48 to V334), when free or bound to BRCA2-HBD. The SEC column is a BIOSEC 3 (Agilent). HSF2BP mutant is dimeric (theoretical mass of the dimer: 64 kDa) and assembles as a tetramer when bound to BRCA2-HBD (theoretical mass of the 4:2 complex: 142 kDa). **b,c** Models of full-length HSF2BP calculated using the program DAMMIF from the SEC-SAXS data. b, Models calculated without making any hypothesis on the symmetry of the HSF2BP oligomer structure. c, Models calculated by hypothesizing that the oligomer structure has a 2-fold symmetry, as it is formed from two dimers. In both cases, 10 models were calculated. The upper panels show the average models. The lower panels display three representative models.

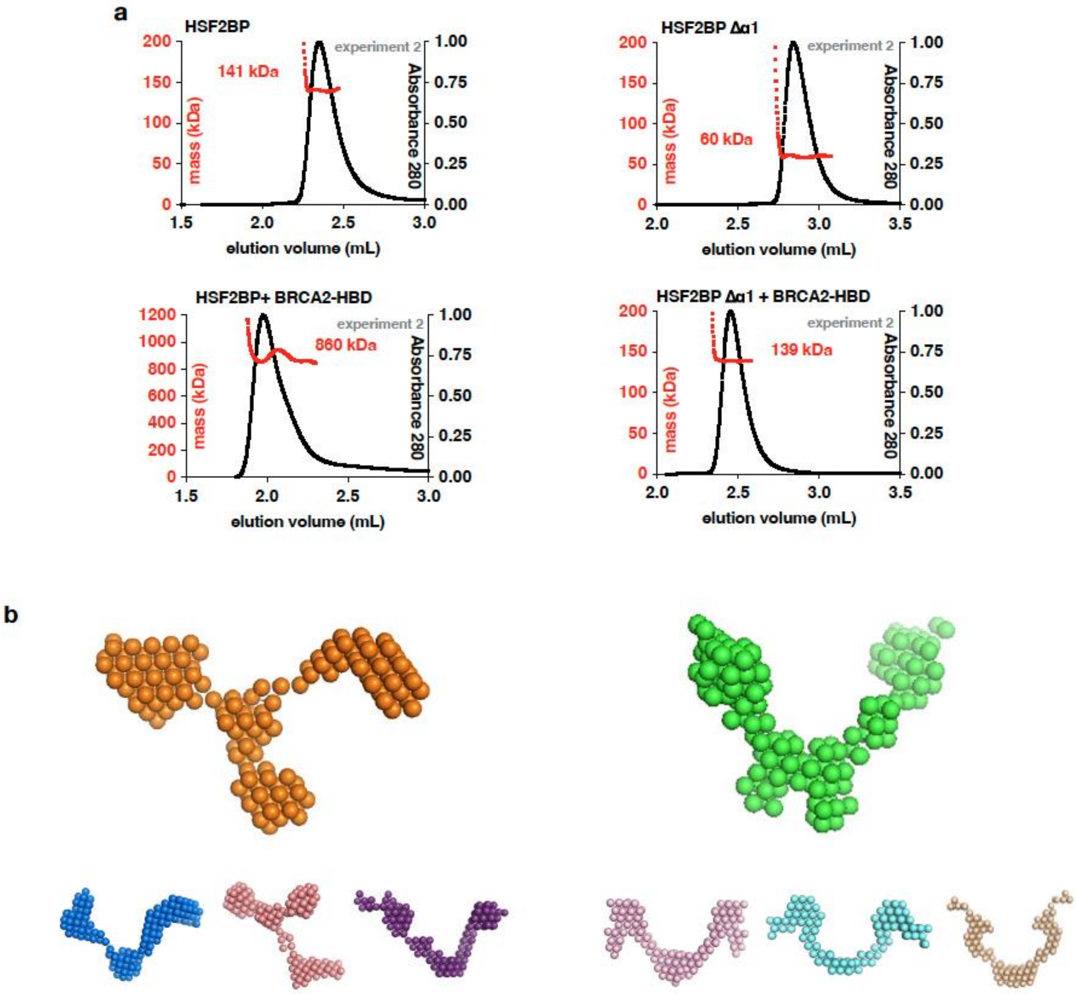

**Extended Data Fig. 2. Extended data for the cryo-EM analysis shown in Fig. 2. a,** A typical 2D class obtained from the analysis of negative-staining images recorded on full-length HSF2BP bound to BRCA2-HBD. **b,** Flow chart of the processing of the cryo-EM data recorded on HSF2BP bound to BRCA2-HBD. The whole analysis was performed using CryoSPARC**.** The first TOPAZ calculation provided more side views of the complex, whereas the second TOPAZ calculation provided more particles. At the end, about 44 particles were picked in average on each micrograph. The best resolved volume was obtained from 8962 x 44,4 = 398 000 particles (see panel **c**). Two other volumes were obtained after 3D classification of the 398 000 particles (see panel **d**).

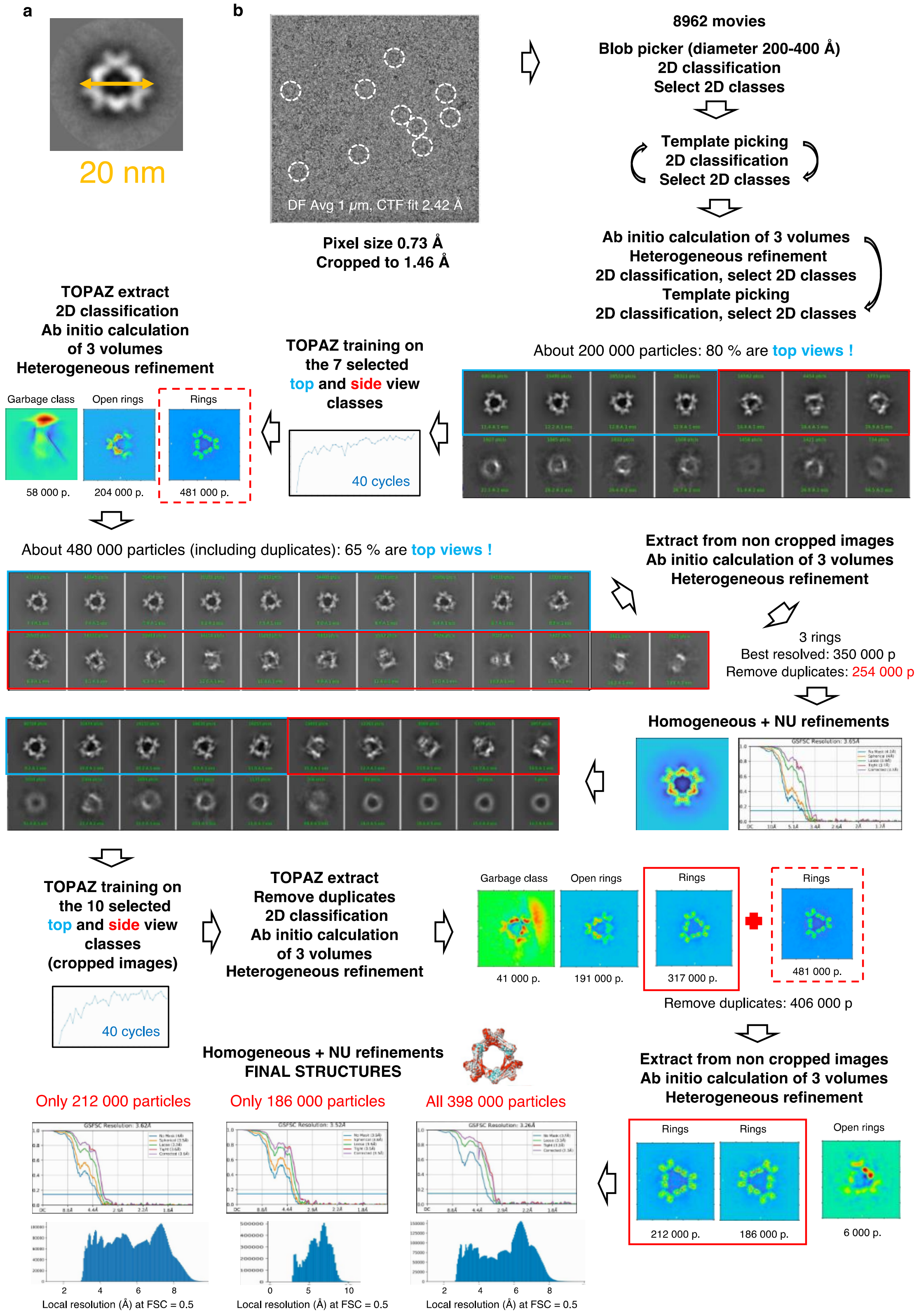
