## Extended data 2c,d-3-4 for "BRCA2-HSF2BP Oligomeric Ring Disassembly by BRME1 Promotes Homologous Recombination"

**c**, Analysis of the final map calculated from 398 000 particles. 2D classes and viewing direction distribution of the particles are presented. Docking of the crystal structure (7BDX) of the complex formed by four armadillo domains of HSF2BP (first and second armadillo dimers in red and orange, respectively) and two BRCA2-HBD peptides (in blue) into the cryo-EM map is illustrated in the lower panels. The armadillo domains are displayed as cartoons, whereas the BRCA2 peptide is displayed in sticks. The map is colored as the fitted crystal structure. The different zoom views highlight that the BRCA2-HBD peptides nicely fit into the cryo-EM map after docking of the HSF2BP armadillo domains. Well-resolved BRCA2 side chains are marked. **d**, Analysis of the cryo-EM maps obtained after classification of the 398 000 particles. The first cryo-EM map, calculated from 212 000 particles (left and grey), has a larger diameter and its resolution is regularly distributed between 3 and 8 Å (see panel **b**). The second cryo-EM map, calculated from 186 000 particles (right and orange), has a slightly smaller diameter; its three inner armadillo domains are better defined, whereas the three outer armadillo domains as well as the coiled-coil regions are poorly resolved.

c

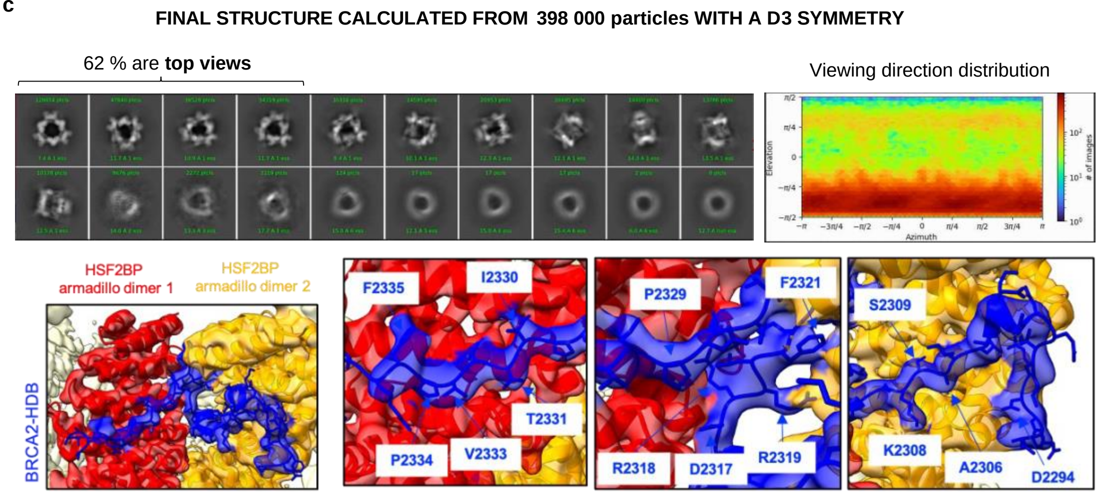


d

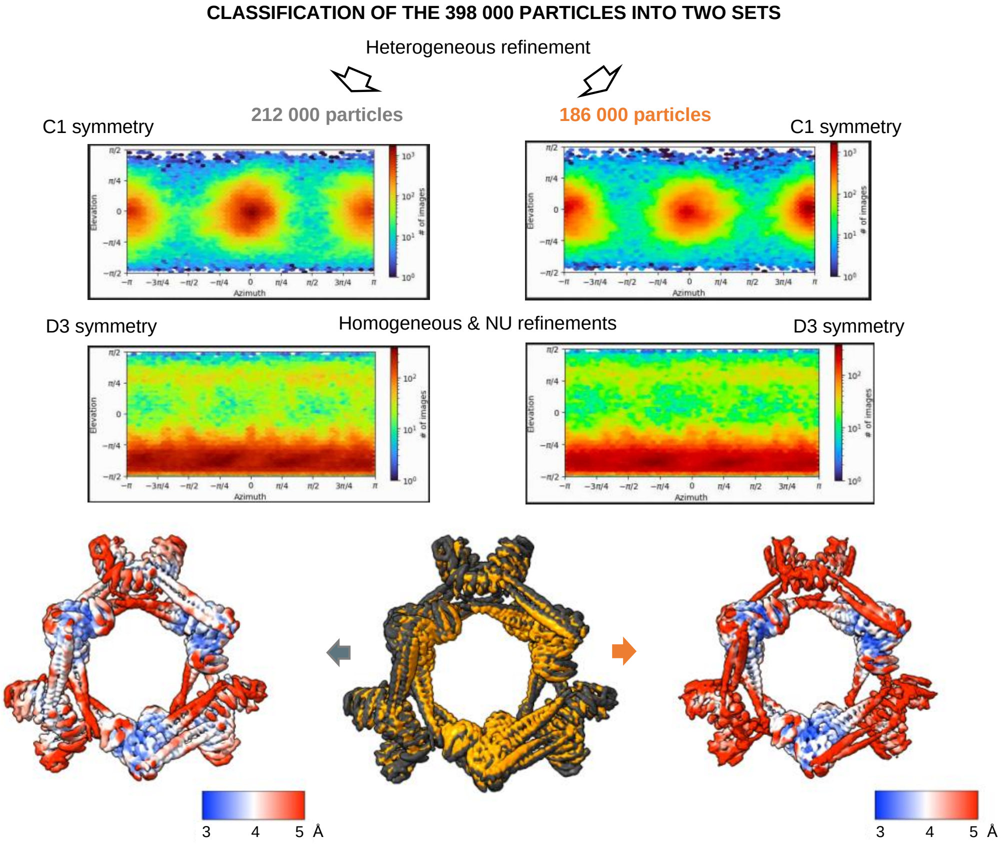

**Extended Data Fig. 3. Extended data for the AlphaFold analysis of the HSF2BP dimer presented in Fig. 3. a**, Five models of the HSF2BP dimer calculated by AlphaFold. In the upper panels are displayed the heat maps showing the predicted relative position error (in Å) calculated by AlphaFold between all pairs of residues (HSF2BP residues from the first and second monomers are numbered as 1-334 and 335-668, respectively). The blue color observed in regions corresponding to intermolecular distances proves that the relative position of the two monomers is predicted with high confidence. In the lower panels, the 5 models are superimposed, each of them being displayed in a different color. They all exhibit the same secondary structure elements. Moreover, their 3D structures are identical, except for the position of the disordered N-terminal region (residues 1 to 19), the β-strand (residues 20 to 24), the first helix α1 (residues 25 to 45) and the loop between α1 and α2 (or hinge; residues 46 to 47), as shown in the main panel and in the zoom view rotated by 90° in the dashed boxed panel. Two putative intermolecular disulfide bridges are displayed in black dots in the main panel. **b**, Superimposition of one of the AlphaFold models of the full-length HSF2BP dimer (in red) onto the crystal structure of the complex between two HSF2BP armadillo dimers (white) and two BRCA2-HBD peptides (blue) (PDB: 7BDX). **c**, Five models of the helix α1 tetramer calculated by AlphaFold**.** The best model is similar to our crystal structure, as shown in the black box (red: AF model; blue: crystal structure; Cα RMSD 0.96 Å). Models 3, 4 and 5 exhibit parallel α-helices, as the α1 tetrameric model that fits into the cryo-EM density (Fig. 3b, inset).

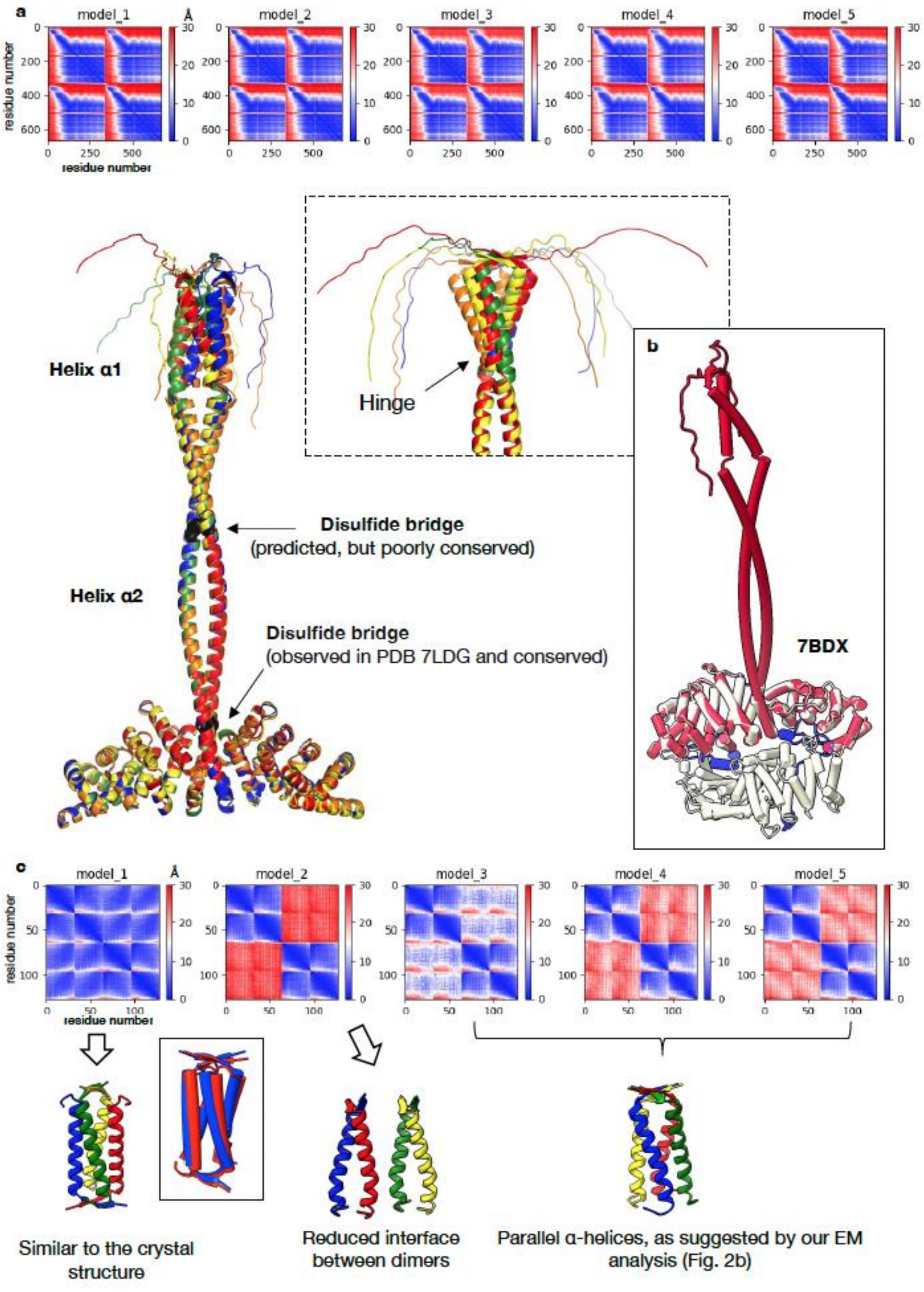

**Extended Data Fig. 4. Extended data for the SEC-MALS analysis presented in Fig. 3.** Replicate of the experiment shown in Fig. 3d and analyses of other HSF2BP variants with substitutions in the residues that appear to be at α2-α2 and α2-armadillo interfaces in the cryo-EM model (Fig. 3c). Proteins were analyzed alone and in complex with BRCA2-HBD, experiments were done twice. Predicted molecular weight of the tetrameric HSF2BP is 150 kDa; the 4:2 complex with BRCA2-HBD — 164 kDa; the 8:4 complex — 328 kDa; the 24:12 complex — 984 kDa.

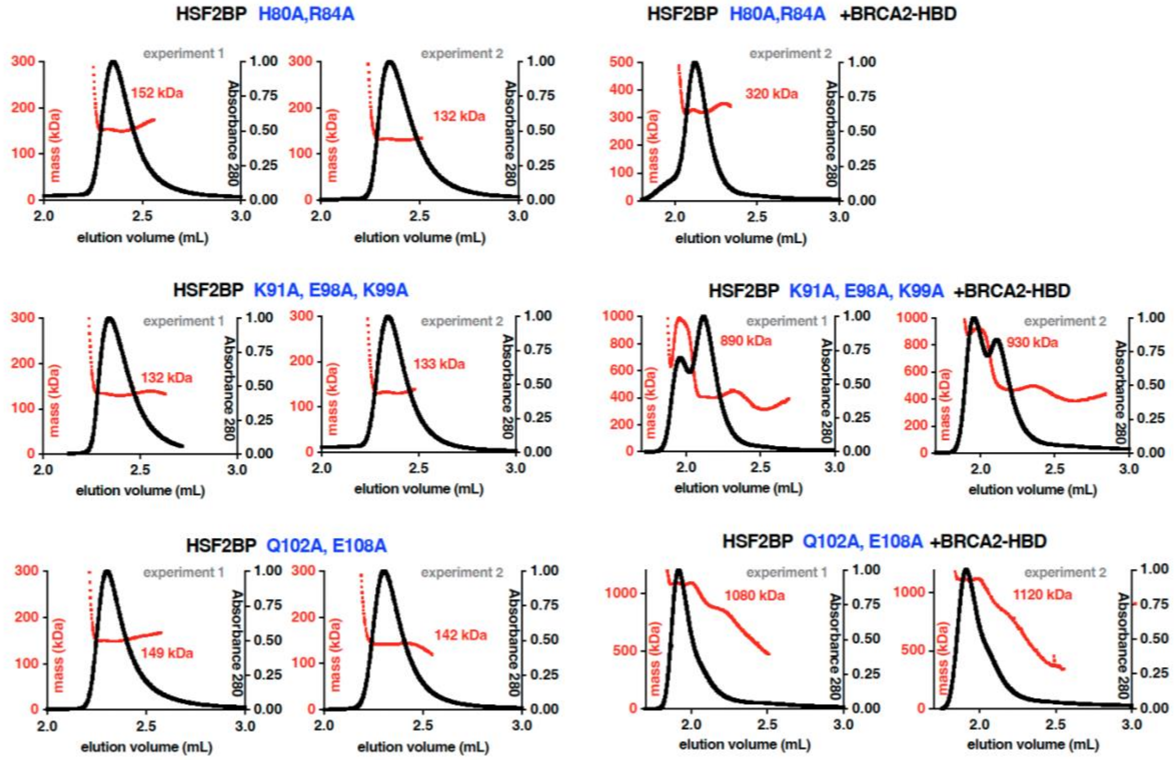
