## Extended data 5-9 for "BRCA2-HSF2BP Oligomeric Ring Disassembly by BRME1 Promotes Homologous Recombination"

**Extended Data Fig. 5. Extended data for the biophysical analyses presented in Fig. 4. a**, Binding of BRME1-M to full-length HSF2BP (left) or HSF2BP helix $\alpha 1$ (right), as analyzed by ITC at 20°C. A large decrease of the binding enthalpy is observed when comparing the affinity of BRME1-M against HSF2BP helix $\alpha 1$ versus full length HSF2BP. Such a decrease is not observed at 30°C, as shown in **Fig. 4b**. All these data are summarized in **Table 1**. **b**, Crystal structure of the $\alpha 1$ tetramer. The HSF2BP peptide E19-V50 is organized as a parallel dimer (each dimer is displayed in red and yellow, respectively), and two of these dimers are interacting in an anti-parallel manner to form the final tetramer. Such conformation was also found by AlphaFold (see **Extended Data Fig. 3c**). Whether it can exist in full-length HSF2BP is yet unclear. However, our cryo-EM analysis strongly suggests that it does not exist in the complex between full-length HSF2BP and BRCA2-HBD.

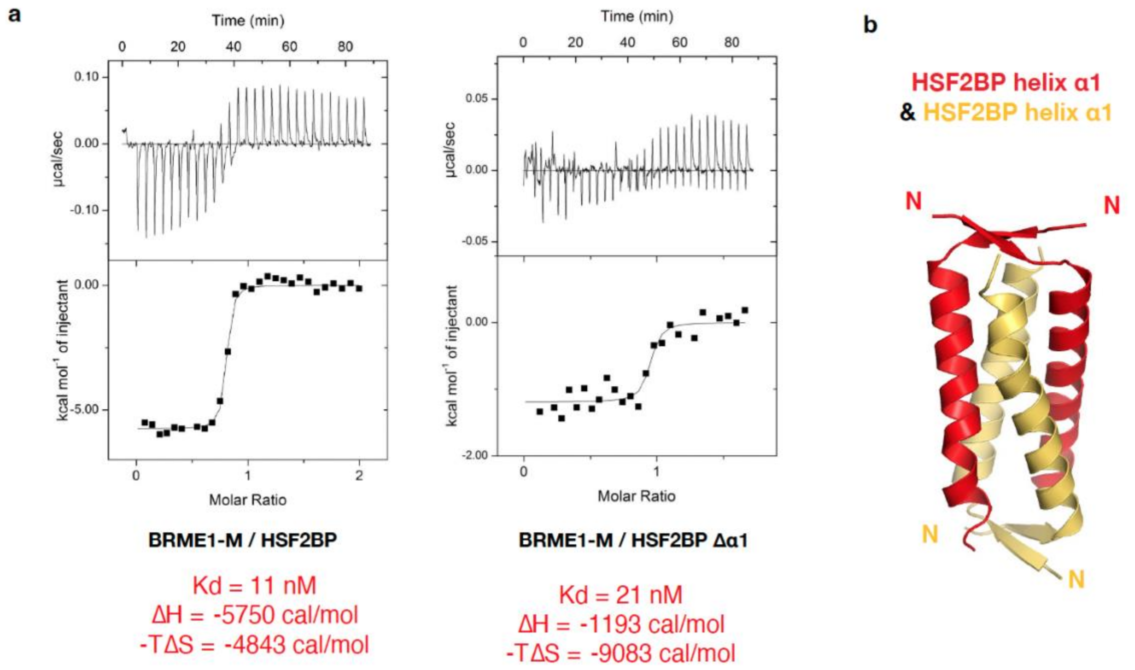

**Extended Data Fig. 6. Replicates for SFM, mass photometry and SEC-MALS experiments shown in Fig. 5. a**,**b** Representative SFM scan images (scale bar = 100 nm) and corresponding density of volumes distributions of the HSF2BP, HSF2BP+BRCA2-HBD and HSF2BP+BRCA2-HBD+BRME1-M complexes. **c**, Mass photometry analysis of free HSF2BP at 25 nM (see Fig. 4e). **d**, Mass photometry analysis of HSF2BP + BRCA2-HBD + BRME1-M to complement experiments shown in Fig. 5e. Top two graphs: replicates of the experiment shown in Fig 5e but at higher concentration (100 nM). Third graph: order of addition was changed by adding BRME1-M 30 min before BRCA2-HBD. Bottom graph: complexes were formed at a lower salt concentration (150 mM NaCl). **e**, SEC-MALS analyses of the HSF2BP-BRME1 complex in the absence (top panel; 1:1 complex theoretical mass: 42 kDa) or in the presence (bottom panel; 2:2:1 complex theoretical mass: 91 kDa) of BRCA2-HBD.

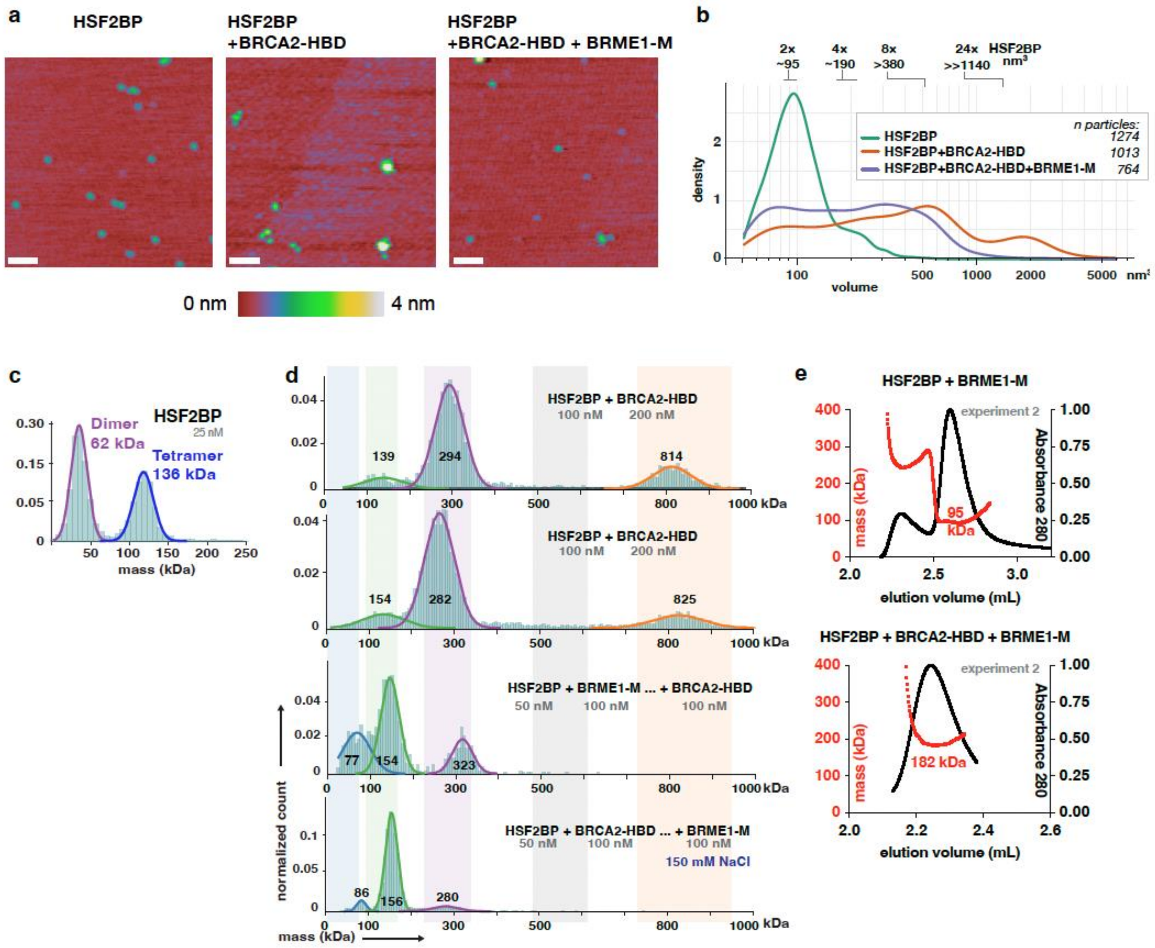

**Extended Data Fig. 7. Extended data for experiments presented in Fig. 6. a**, Anti-GFP immunoblot of the total protein extract from HeLa cells stably producing GFP-HSF2BP variants or GFP. **b**, Clonogenic survivals of HeLa cells stably producing the indicated GFP-HSF2BP variants or GFP. Surviving fraction after treatment with the indicated concentrations of mitomycin C (MMC), cisplatin or talazoparib is plotted on a log scale**.** Each experiment was repeated three times, means and s.e.m. are displayed. **c,d**, Replicates of the cisplatin crosslink repair assay shown in Fig. 6f. **e,** Replicate of the 2D gel electrophoresis of HR intermediates shown in Fig. 6g. **f**, Replicate of the immunoblot experiment shown in Fig. 6h.

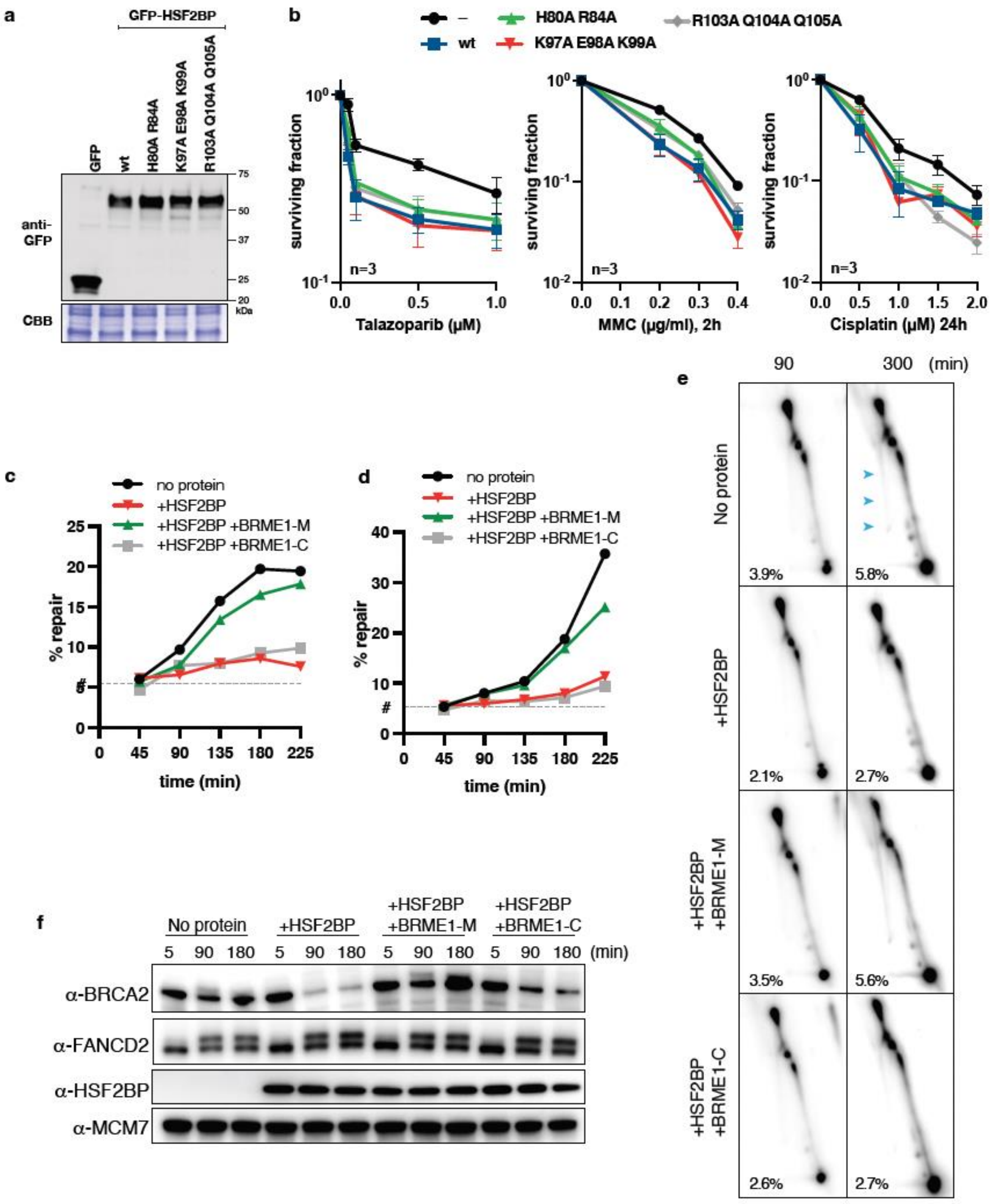

**Extended Data Fig. 8. Analysis of the surface properties of HSF2BP. a**, Different views of the surface of the HSF2BP dimer, as modeled by AlphaFold. This surface is colored as a function of the conservation of the residues in HSF2BP homologs, from red (non-conserved) to blue (strictly conserved) (conservation scores calculated using Consurf: https://consurf.tau.ac.il [58]). Three conserved patches are identified: one patch on helix $\alpha$1; one patch on the armadillo domain and helix $\alpha$2; one patch on the armadillo domain, at the interface with BRCA2. **b**, Interaction of one HSF2BP dimer, displayed as in panel **a**, with two other HSF2BP dimers, displayed in dark grey, in the HSF2BP-BRCA2 model. Interdimeric contacts are observed between $\alpha$1 dimers, an $\alpha$2 coiled coil and an armadillo domain, and two $\alpha$2 coiled coils. These interfaces are at least partially conserved. **c**, Analysis of the electrostatic potential at the surface of a 3D model of the complex formed by 24 HSF2BP molecules, as shown in **Fig. 3a.** This surface is colored from red (negatively charged) to blue (positively charged). It is mainly negatively charged. All these images were produced using Pymol 2.5.2.

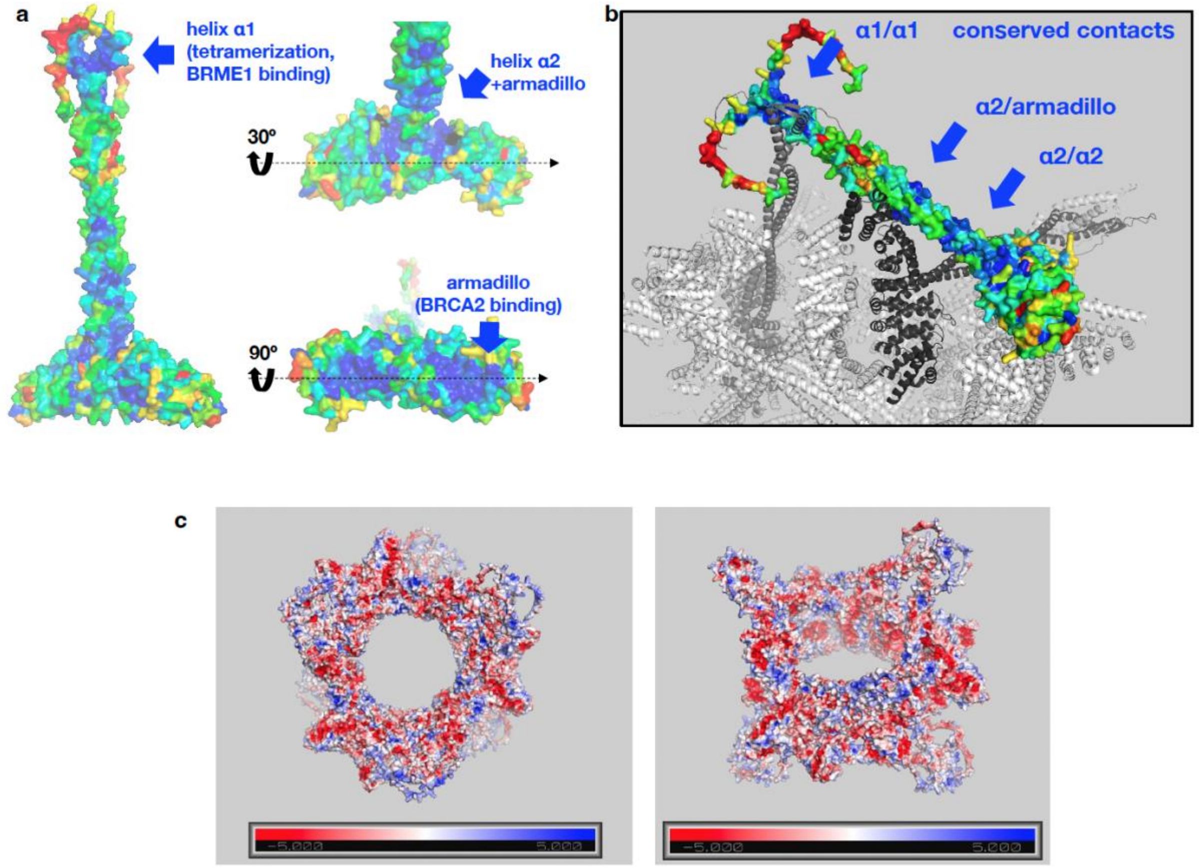

**Extended Data Fig. 9. Cysteine residues contribute to HSF2BP function. a**, Sequence alignment of human HSF2BP with ten homologs from mammals to fishes, mollusk and marine worm. Positions corresponding to human HSF2BP cysteines are boxed. **b,** Position of the cysteines in the 3D structure of dimeric HSF2BP. The armadillo dimer is shown in the main panel (PDB 7LDG), and part of the helix $\alpha 2$ coiled coil is displayed in the black box (AlphaFold model). The BRCA2-HBD peptide is in yellow. The cysteines are in purple. **c**, SDS-PAGE gels of the GFP-HSF2BP variants expressed in HeLa cells. **d**, Survival assays performed with cells expressing the single cysteine mutants C78A, C120S and C128A, as well as the triple mutant combining the three mutations.

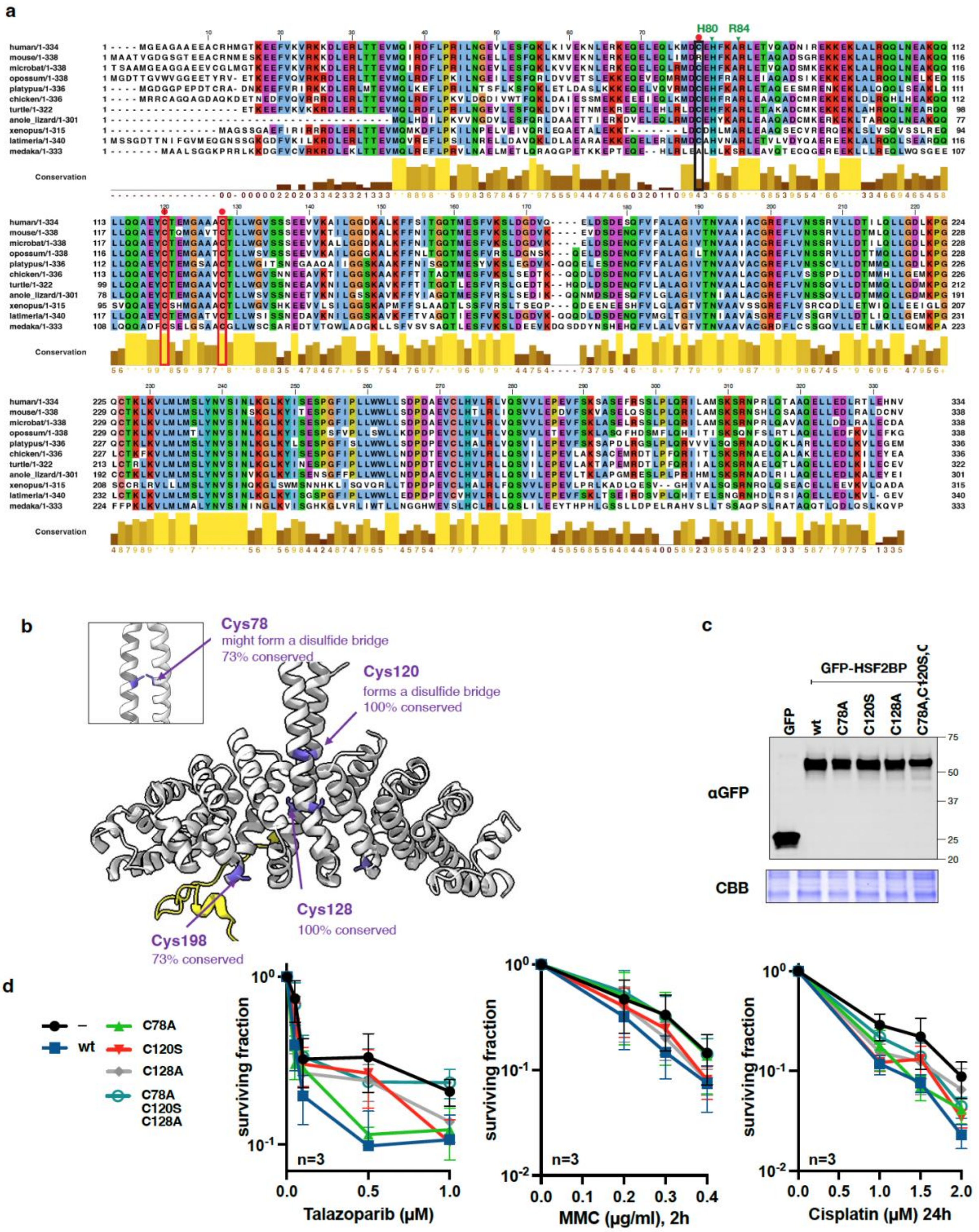
